# Ribosome collisions trigger tmRNA-mediated rescue through mRNA disengagement

**DOI:** 10.64898/2026.04.23.720132

**Authors:** Mikhail Metelev, Anneli Borg, Shirin Akbar, Daniel S. D. Larsson, A. Carolin Seefeldt, Maria Selmer, Magnus Johansson

**Author notes:** These authors contributed equally (Mikhail Metelev, Anneli Borg).

## Abstract

Ribosome rescue pathways maintain translation during normal growth and contribute to stress tolerance. In *Escherichia coli*, HrpA is a ribosome-collision recognition factor that mediates antibiotic tolerance by resolving stalled ribosomes, yet the fate of the resulting ribosomal intermediate, with a topologically trapped peptidyl-tRNA in the 50S nascent peptide exit tunnel, has remained unclear. Using *in vivo* single-molecule tracking, we show that under antibiotic treatment, HrpA-dependent collision processing promotes recruitment of tmRNA and Hsp15 as downstream rescue factors. Cryo-electron microscopy of affinity-purified native complexes from tmRNA-deficient cells reveals accumulation of aberrant 70S·peptidyl-tRNA complexes and shows that Hsp15, previously thought to interact exclusively with 50S·peptidyl-tRNA, binds to 70S complexes lacking mRNA in the E site. This intermediate can be resolved by the canonical tmRNA pathway, thereby connecting HrpA-dependent collision processing to canonical tmRNA-mediated rescue.

## INTRODUCTION

Translating ribosomes often stall on truncated, damaged, or difficult-to-decode mRNAs, producing either non-stop complexes at the 3′ end of a transcript or no-go stalls caused by internal roadblocks^1,2^. If left unresolved, stalled ribosomes accumulate in inactive complexes, which rapidly diminish the cell’s ability to synthesize proteins. Bacteria rely on dedicated rescue pathways to return these ribosomes to the active pool, the most widespread of which is trans-translation, the principal pathway for resolving non-stop complexes (Fig. 1a)^3^.

**Fig. 1.**
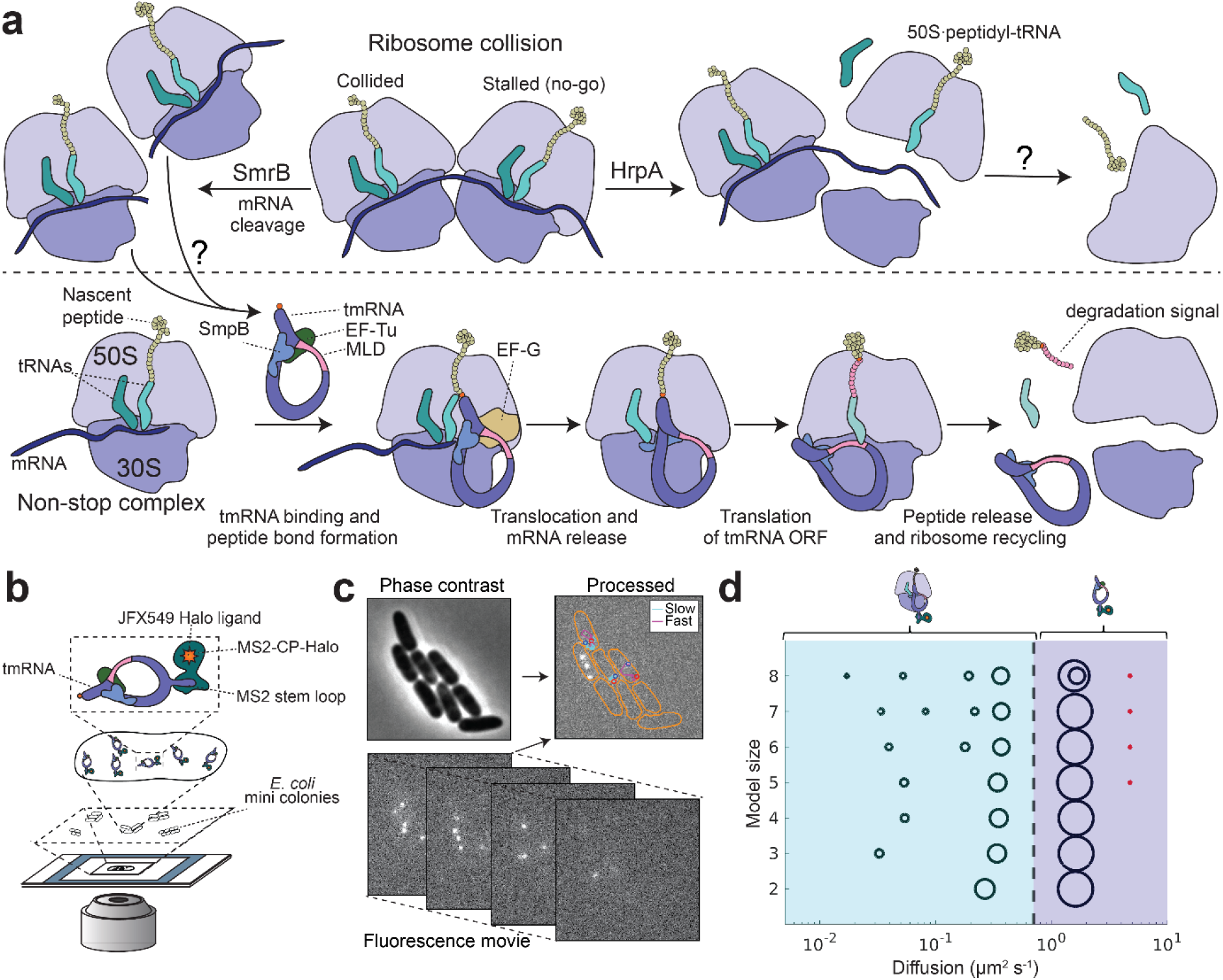
Overview of aberrant ribosome processing in *E. coli* and experimental design. **a**. Schematic overview of ribosome surveillance pathways, showing ribosome collision processing by SmrB and HrpA (top), tmRNA-dependent rescue of non-stop ribosomes (bottom), and the proposed link between them. **b**. Fluorescence labeling strategy (illustrated for tmRNA) and single-molecule tracking microscopy setup. **c**. Phase contrast and fluorescence time-lapse imaging of labelled cells used for segmentation and single-molecule trajectory building. Cell outlines are shown in orange; trajectory start and end points are indicated in red and blue, respectively. Trajectories are color-coded by diffusive state. The figure shows diffusion trajectories of tmRNA-MS2^g^ (see also Supplementary Movie 1). **d**. HMM fitting results of single-molecule trajectories of tmRNA-MS2^g^ using models with 2–8 states (Supplementary Data 1). Discrete diffusion states are shown as circles positioned according to their diffusion coefficients. The area of each circle represents the relative occupancy of the corresponding diffusional state. The black dashed line indicates the threshold at 0.7 µm^2^ s^−1^ separating ribosome-bound states (blue) from free tmRNA-MS2 (violet). The figure shows the results from HMM fitting of combined datasets from seven individual experiments

The trans-translation system consists of a specialized hybrid RNA molecule, tmRNA, which contains two functional modules: a tRNA-like domain (TLD) and an mRNA-like domain (MLD)^4,5^. The TLD is aminoacylated with alanine and delivered to the ribosome by EF-Tu^6^. Together with its partner SmpB, it mimics a tRNA and recognizes non-stop ribosomes whose A-site mRNA channel is empty or contains only a short 3’ mRNA extension, the hallmark of non-stop complexes^7^. After accommodation, the nascent peptide is transferred from the P-site tRNA to Ala-tmRNA, the obstructing mRNA is released, and translation resumes on the MLD, which encodes a short alanine-rich peptide tag that targets the incomplete polypeptide for degradation. Many bacteria carry alternative rescue systems such as ArfA and ArfB^8,9^. In *E. coli*, loss of either trans-translation or ArfA alone has only a minor phenotypic effect, but deleting both is synthetically lethal, highlighting the essential role of ribosome rescue^8^.

In addition to rescuing non-stop complexes, tmRNA can also target ribosomes stalled on internal mRNA sequences that induce prolonged pausing, such as stretches of rare codons ^10^. The prevalent model suggests that these no-go stalls are first processed by mRNA cleavage, which converts them into non-stop substrates for trans-translation^11^. Recent studies reveal that, as in eukaryotes, ribosome collisions arising from prolonged pausing can serve as a universal signal for initiating rescue pathways in bacteria^12,13^. Different bacteria have independently evolved diverse mechanisms to act downstream of collision sensing^14–16^.

In *E. coli*, two collision-specific factors have recently been identified (Fig 1a)^14,16^. The small nuclease SmrB binds the interface between a stalled ribosome and its trailing collided partner and cleaves the mRNA between them^16^. This activity converts the collided ribosome into a non-stop complex that can, according to the suggested model, be rescued by tmRNA. Rescue of the leading stalled ribosome is thought to rely on mRNA degradation by additional nucleases. Notably, deletion of *smrB* confers no phenotypic cost, even under antibiotic-induced stalling, and its physiological role therefore remains to be fully defined^14^.

A second factor, the large ATP-dependent RNA helicase HrpA, has been implicated in tolerance to ribosome-targeting antibiotics^14^. HrpA recognizes the interface between two collided ribosomes and requires downstream mRNA for its activity. *In vitro* reconstitution experiments indicate that HrpA can split the leading stalled ribosome into subunits, consistent with a model in which helicase engagement with downstream mRNA exerts a pulling force, destabilizing the stalled ribosome^14^. *In vivo*, however, HrpA activity leads to the accumulation of 70S monosomes, which has been interpreted as a possible reassociation of subunits after an initial splitting step, releasing the mRNA^14^. Despite recent progress, key steps in how HrpA remodels collided ribosomes and facilitates rescue remain unclear. In particular, if splitting does occur, it results in the generation of either 50S subunits bearing peptidyl-tRNA (50S·peptidyl-tRNA) or, possibly, reassociated 70S complexes with peptidyl-tRNA but no mRNA (70S·peptidyl-tRNA). How such complexes are processed in *E. coli* remains unresolved.

Many bacteria possess a ribosome quality control (RQC) pathway, mediated by RqcH and RqcP, that act on 50S·peptidyl-tRNA complexes to promote tagging and degradation of aberrant nascent peptides^17–19^. In this pathway, RqcH recruits alanine-charged tRNAs to extend the stalled nascent chain, whereas RqcP stabilizes and orients the peptidyl-tRNA on the 50S subunit to support this reaction in the absence of the 30S. The resulting elongated nascent chain is ultimately released by peptidyl-tRNA hydrolase (Pth) and targeted for degradation^20^. This is conceptually similar to the eukaryotic pathway, in which 60S·peptidyl-tRNA complexes are recognized by Rqc2/NEMF, leading to recruitment of the E3 ligase Ltn1/Listerin, which ubiquitinates the nascent chain and targets it for proteasome-mediated degradation^21–24^.

How 50S·peptidyl-tRNA complexes are processed in *E. coli* remains unresolved. *E. coli* lacks RqcH and instead encodes only a homolog of RqcP, Hsp15, a heat-shock–induced factor that binds such complexes with high affinity in a manner similar to RqcP^25^. Notably, Hsp15 contains a long C-terminal α-helix that is absent in species where its homolog functions together with RqcH. Treatment of *E. coli* with chloramphenicol has been shown to promote Hsp15 binding to aberrant 50S subunits^26^; however, whether Hsp15 functions in ribosome rescue in addition to its role in the heat-shock response has remained unknown. Interestingly, tmRNA mutants are sensitive to heat stress, and tmRNA activity increases at elevated temperatures^27^, likely reflecting increased ribosome stalling. However, how tmRNA gains access to these complexes is not clear.

Here we use single-molecule tracking to characterize the ribosome-binding dynamics of tmRNA, SmrB, HrpA, and Hsp15 in live cells under normal growth and under erythromycin-induced stress, triggering ribosome collisions. By quantifying their recruitment to ribosomes, we discover an unexpected link between tmRNA and HrpA, indicating that tmRNA acts downstream of HrpA, contrary to the current model. Furthermore, we show that Hsp15 also functions downstream of HrpA and that Hsp15-bound ribosomes are preferentially processed by tmRNA, consistent with the formation of a shared 70S intermediate. Using cryo-electron microscopy (cryo-EM) and cross-linking, we show that Hsp15 binds not only 50S·peptidyl-tRNA complexes but is also present on 70S·peptidyl-tRNA complexes, and that its C-terminal α-helix extends into the empty E-site mRNA channel on the 30S subunit. Together, these findings reveal an intricate network of cooperation among bacterial ribosome rescue factors, and suggest a solution to the long-standing problem of a missing RQC pathway in *E. coli*.

## RESULTS

### Tracking of tmRNA, HrpA, and SmrB

To fluorescently label tmRNA for single-molecule tracking in live *E. coli*, we used an approach similar to that previously developed for tracking of ribosomal subunits^28^. An MS2 stem-loop aptamer was introduced into the genomic *ssrA* gene, which encodes tmRNA (Fig. 1**b**, Supplementary Fig. 1a). Whereas deletion of *ssrA* caused a growth defect and increased sensitivity to erythromycin, the MS2-tagged *ssrA* strain (tmRNA-MS2^g^) displayed a doubling time similar to the WT strain (Supplementary Fig. 2), showing that the modified tmRNA remained functional. Moreover, *arfA* could be deleted in the MS2-tagged *ssrA* background, in contrast to the known synthetic lethality of the *ssrA* and *arfA* double deletion. Low-level expression of an MS2-coat protein fused to a HaloTag protein (MS2CP-HaloTag) allowed labeling of a subpopulation of tmRNA molecules in the tmRNA-MS2^g^ strain.

HrpA and SmrB were fluorescently labeled by N-terminal HaloTag fusions. Available structures indicate that the N terminus of HrpA is not involved in the interaction with the collided ribosomes, and that SmrB, which binds largely in between the collided ribosomes, has its N terminus exposed and oriented away from the ribosomal collision interface (Supplementary Fig. 1b-c).

For all single-molecule tracking experiments we used an approach previously established to track fluorescently labeled ribosomal subunits and factors involved in translation (Fig. 1b-c) ^28–32^. In this approach, HaloTag fusions were covalently labeled inside cells with a JFX549 HaloTag ligand. The cells were subsequently grown on agarose pads containing rich defined medium (RDM) to form mini-colonies, resulting in even dilution of fluorescently labeled molecules among progeny cells. For each labeled factor, more than 1,000 individual cells were imaged using stroboscopic laser illumination (3 ms) at a 30 ms camera exposure, and corresponding phase-contrast images were collected for cell segmentation. We observed complex diffusional behavior for the tracked factors with populations of slow- and fast-moving molecules consistent with recruitment of the factors to the ribosome (Supplementary Movie 1-3, top panel).

The fluorescence movies were analyzed using a semi-automatic analysis pipeline in which cells were segmented, fluorescent spots were detected and linked across frames to generate single-molecule diffusion trajectories (Fig. 1, Supplementary Movie 1-3). Average trajectory lengths ranged from 11 to 17 frames for the different factors, with some trajectories longer than 100 frames. Ensembles of trajectories from individual microscopy experiments were analyzed using Hidden Markov modeling (HMM) to infer state-specific diffusion coefficients, occupancies, and dwell times calculated from frequencies of transitions between diffusion states, with the number of states defined a priori^32,33^. For each of the labeled molecules we fit model sizes between 2 and 8 states and compared the results with those obtained from 50S subunit tracking data performed as described previuosly^28^ (Supplementary Fig. 3, Supplementary Data 1-4). Ribosomes exhibit a broad range of diffusive states, likely corresponding to free ribosomal subunits, polysomes, and ribosomes engaged in higher-order macromolecular complexes, including those associated with RNA polymerase or the cell membrane (Supplementary Fig. 3, Supplementary Data 4). With increasing HMM model size, all tracked factors display clusters of diffusive states similar to those observed for ribosomes, as well as faster diffusing states likely corresponding to free diffusion (Fig. 1d, Supplementary Fig. 3). To distinguish free and ribosome-bound states we introduced a threshold at 0.7 µm^2^ s^−1^ which we used for coarse graining of models with ≥ 3 states into two-state diffusion models (Supplementary Fig. 3, Supplementary Data 1-4). We found that diffusion coefficients, steady-state occupancies and dwell times were robust regardless of the underlying HMM model size used for the coarse graining and reproducible across independent microscopy experiments (Supplementary Fig. 4, Supplementary Data 1-3). As coarse-graining of larger HMM models yielded comparable results, we therefore decided to use a three-state HMM model coarse-grained into two diffusive states to capture ribosome-binding kinetics for all subsequent analyses, unless otherwise specified. For a discussion regarding the use of coarse-graining, rather than solely relying on 2-state HMM fittings, see Supplementary note 1.

Our previous ribosome tracking study showed that free ribosomal subunits diffuse at ∼0.5 µm^2^ s^−1^, whereas mRNA-bound ribosomes display slower and heterogeneous diffusion (0.001–0.25 µm^2^ s^−1^)^28^. tmRNA-MS2^g^ exhibited an average diffusion coefficient of 0.28 ± 0.03 µm^2^ s^−1^ in the ribosome-bound state, intermediate between free subunits and mRNA-bound 70S ribosomes (Supplementary Data 1). Analysis of HMM models with ≥3 states resolved two ribosome-associated substates: a predominant faster diffusing substate (0.34 ± 0.02 µm^2^ s^−1^), showing modest nucleoid exclusion, and a less populated substate with approximately an order-of-magnitude slower diffusion (0.03 ± 0.01 µm^2^ s^−1^), enriched at the cell periphery (Supplementary Fig. 5, Supplementary Data 1). These observations suggest that tmRNA-MS2^g^ is predominantly observed in a state corresponding to individually diffusing 70S ribosomes engaged with the short tmRNA open reading frame (ORF). Such 70S monosomes are expected to exhibit nucleoid exclusion driven by nucleoid structure, geometrical confinement and macromolecular crowding^34^ in agreement with our observations. However, since tmRNA resumes the elongation of nascent polypeptides on stalled ribosomes, a fraction of these is expected to already be membrane-inserted at translocons, giving rise to a subcluster with membrane localization, in line with our observations.

tmRNA-MS2^g^ occupies the ribosome-associated state with a steady-state occupancy of 25 ± 0.3% and a mean dwell time of 3.6 ± 0.7 s (Supplementary Data 1). This shows that tmRNA– ribosome complexes persist for several seconds once formed. Since the fitted diffusion rate of the highly populated cytoplasmic ribosome-bound state corresponds well with the expected diffusion of single 70S particles rather than polysomes, our result suggests that translation of the internal tmRNA ORF determines the lifetime of the ribosome-associated state. The initial steps of rescue, including association with non-stop ribosomal complexes and mRNA release, are either rapid and therefore not resolved by our analysis, or occur at low frequency.

We also performed single-molecule tracking of previously characterized trans-translation mutants using the tmRNA-MS2^g^ system (Supplementary Fig. 6, Supplementary Data 5-7). Firstly, in the absence of its partner SmpB, tmRNA-MS2^g^ showed a marked decrease in slow-state occupancy, from 25 ± 3% to 6 ± 2%. (Supplementary Fig. 6, Supplementary Data 5). A similar decrease was observed for an *ssrA* G3A acceptor stem mutant of tmRNA, which is known to impair alanine charging and thereby EF-Tu-dependent delivery of tmRNA to the ribosome (Supplementary Fig. 6, Supplementary Data 6)^6,35^. In addition, we tested an *ssrA* A86C mutant, which has been suggested to bind to stalled ribosomes and accept the nascent peptide from the P-site tRNA but is then unable to proceed with translation of the tmRNA-encoded ORF, presumably resulting in ribosome stalling^36,37^. Consistent with this, we observed a marked increase in ribosome-bound occupancy to 92 ± 2% (Supplementary Fig. 6, Supplementary Data 7). Taken together, these results indicate that the tmRNA-MS2^g^ variant is functional and that the coarse-graining threshold allows us to correctly distinguish free and ribosome-bound states.

We observed that HaloTag-HrpA occupies the ribosome-bound state with an occupancy of 35 ± 4%, a dwell time of 1.0 ± 0.1 s, and a diffusion coefficient of 0.06 ± 0.01 µm^2^ s^−1^ (Supplementary Data 2). The substantially slower diffusion compared to tmRNA-MS2^g^ is consistent with HrpA binding to stalled ribosomes in the polysomal fraction that diffuse more slowly than an mRNA-free 70S complex. For HaloTag-SmrB, in HMM models with ≥ 3 states the slow diffusion states divide in two subclusters: one major, occupying states similar to those observed for HaloTag-HrpA (16% occupancy), and one minor with a diffusion coefficient of ∼0.5 µm^2^ s^−1^ (5% occupancy) (Supplementary Data 3). Both HrpA and SmrB displayed a stronger tendency toward nucleoid exclusion in slow diffusion states than tmRNA (Supplementary Fig. 5) consistent with the expectation that polysomes, with which these factors are expected to associate, exhibit nucleoid exclusion^34^.

We next tested whether deletion of either of the collision-sensing factors leads to increased ribosome engagement of the other. Deletion of *smrB* had no detectable effect on the extent of HaloTag-HrpA association to ribosomes (Supplementary Fig. 7, Supplementary Data 8). In contrast, deletion of *hrpA* resulted in an increase in the ribosome-associated fraction of HaloTag-SmrB, suggesting that substrates normally processed by HrpA are redirected to the SmrB pathway (Supplementary Fig. 7, Supplementary Data 9). When HaloTag–HrpA and HaloTag–SmrB were tracked in a double-deletion *ΔsmrBΔhrpA* strain, both showed increased ribosome-bound state occupancy, suggesting accumulation of collided ribosomal complexes in the absence of the endogenous rescue factors (Supplementary Fig. 7, Supplementary Data 10-11).

Previous studies have shown that, even under normal growth conditions, ribosomes frequently fail to complete translation and require rescue by tmRNA^38^. Our results support this view and also indicate that ribosome collisions commonly occur under normal growth as we observe frequent ribosome engagement by HrpA and SmrB.

### Tracking of tmRNA, HrpA, and SmrB under erythromycin treatment

To investigate how tmRNA, HrpA and SmrB participate in ribosome rescue under stress conditions, we performed single-molecule tracking of these labeled factors in cells treated with a subinhibitory concentration of erythromycin (25 µg ml^−1^), at which wild-type cells show 30% increase in doubling time (Supplementary Fig. 2). Erythromycin has been shown to stall elongating ribosomes at specific peptide sequences, leading to ribosome collisions^14,16,39,40^.

Upon erythromycin treatment, all factors show more than a twofold increase in the occupancy of ribosome-associated states, reaching 66 ± 2%, 73 ± 2%, and 47 ± 8%, for tmRNA-MS2^g^, HaloTag-HrpA, and HaloTag-SmrB, respectively (Fig. 2a, Supplementary Data 12-14). Analysis of the spatial distributions revealed moderate nucleoid exclusion of the ribosome-bound fraction of tmRNA–MS2^g^ (Fig. 2b). The slow diffusion state can still be resolved into two slow diffusive substates: a less populated very slow diffusion (0.05 ± 0.01 µm^2^ s^−1^) state with membrane localization and a highly populated state with a diffusion coefficient of 0.28 ± 0.02 µm^2^ s^−1^ which shows cytosolic localization with moderate nucleoid exclusion (Fig. 2b, Supplementary Fig. 8, Supplementary Data 12). In contrast, ribosome-associated HaloTag-HrpA and HaloTag-SmrB predominantly occupy very slow diffusive states, with an average diffusion coefficient of 0.07 µm^2^ s^−1^ and 0.05 µm^2^ s^−1^ (Supplementary Data 13-14), respectively, and exhibit pronounced nucleoid exclusion (Fig. 2b). These results are consistent with a model in which SmrB and HrpA are recruited to slowly diffusing ribosomes within polysomes, where collisions are expected to occur, whereas tmRNA is predominantly associated with single 70S ribosomes, as reflected by the differences in their diffusion coefficients. Together, these observations demonstrate that erythromycin induces aberrant translation that is sensed by these quality-control factors.

**Fig. 2.**
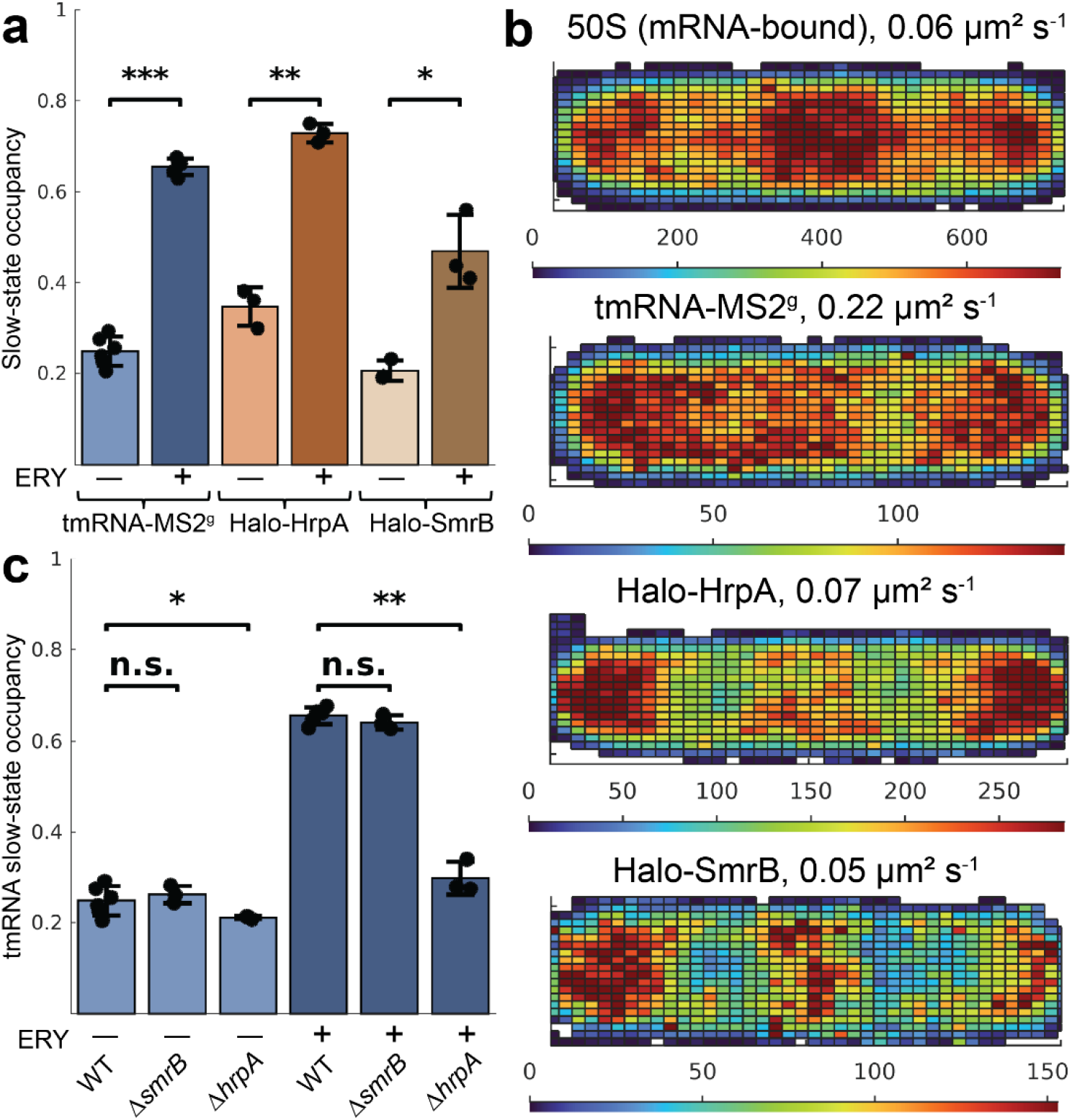
Recruitment of tmRNA-MS2^g^, HaloTag-SmrB and HaloTag-HrpA for ribosome rescue during erythromycin treatment. **a**. Ribosome-bound state occupancies in untreated and erythromycin-treated cells. **b**. Spatial distributions of mRNA-bound 50S subunits and ribosome-bound factors in erythromycin-treated cells. The average diffusion coefficients in those states are indicated. **c**. Ribosome-bound state occupancies of tmRNA-MS2^g^ in WT, *ΔsmrB* and *ΔhrpA* strains without or with erythromycin treatment. Bars represent averages from independent experiments, and error bars indicate standard deviations between experiments. Statistical significance between groups was assessed using a two-sided unpaired t-test. P-values are indicated in the figure as follows: P < 0.05 (*), P < 0.01 (**), P < 0.001 (***), and not significant (n.s.) otherwise.

### tmRNA recruitment under antibiotic stress is linked to HrpA but not SmrB

It has recently been proposed that SmrB-mediated mRNA cleavage targets collided ribosomes for tmRNA-dependent rescue, whereas HrpA activity promotes splitting of stalled 70S ribosomes into 30S and 50S·peptidyl-tRNA which are subsequently resolved by an as-yet unknown mechanism in *E. coli*^*14,16*^. To test whether the increased SmrB activity under erythromycin treatment (Supplementary Fig. 7) constitutes a major pathway for generating substrates for tmRNA, we tracked tmRNA-MS2^g^ in a Δ*smrB* strain. In parallel, we tracked tmRNA-MS2^g^ in a Δ*hrpA* strain, for which an increase in tmRNA recruitment was expected if a larger fraction of ribosome collisions is processed through the SmrB pathway (Supplementary Fig. 7). Surprisingly, the deletion of *smrB* had no detectable effect on tmRNA-MS2^g^ recruitment, neither under non-stress conditions nor under erythromycin treatment (Fig. 2c, Supplementary Data 1,12,15,17). Strikingly, however, the deletion of *hrpA* resulted in a marked decrease in the fraction of tmRNA-MS2^g^ associated with ribosomes under erythromycin treatment, and a smaller decrease under untreated conditions (Fig. 2c, Supplementary Data 1,12,16,18). Thus, both these results are in stark contrast to the proposed mechanism of action of HrpA and SmrB (Fig 1a). Instead of the proposed connection between SmrB and tmRNA it appears as if HrpA action is producing ribosome complexes that need subsequent processing by tmRNA. To test whether the observed effects could be explained by differences in tmRNA expression levels, we quantified the tmRNA-MS2^g^ abundance in otherwise wild-type, *ΔhrpA*, and *ΔsmrB* backgrounds under non-stress conditions and during erythromycin treatment. We found that the observed differences in occupancy could not be explained by differences in tmRNA-MS2^g^ abundance in those strains (Supplementary Fig. 9).

In *E. coli*, in addition to tmRNA, two proteins, ArfA and ArfB, are known to rescue non-stop stalled ribosomes, with ArfA expression being directly regulated by tmRNA^8,41,42^. To exclude the possibility that these alternative pathways contribute to the observed effects in the *ΔsmrB* and *ΔhrpA* strains, we decided to track tmRNA in the corresponding strains carrying additional deletions of both *arfA* and *arfB*. For this purpose, we created a plasmid enabling low-level expression of labelled tmRNA (tmRNA-MS2^p^). First, we used this plasmid-based tmRNA-MS2^p^ system to perform tracking experiments in strains carrying deletions of either *hrpA* or *smrB* (Supplementary Fig. 10a, Supplementary Data 19-21). We noted that plasmid-based tmRNA tracking resulted in an overall lower fraction of tmRNA in the ribosome-associated state, despite lower total tmRNA levels compared with the genomically tagged strain (Supplementary Fig. 11), likely reflecting more efficient clearance of stalled ribosomes by endogenous wild-type tmRNA than by tmRNA-MS2. Nevertheless, under erythromycin treatment, tmRNA activity remained linked to HrpA (Supplementary Fig. 10a, Supplementary Data 22-24). We then performed tracking in strains in addition lacking both *arfA* and *arfB* and, notably, even then, the tmRNA activity remained linked to HrpA but not to SmrB (Supplementary Fig. 10b, Supplementary Data 25-30).

In summary, our results show that under antibiotic treatment, tmRNA is recruited to stalled ribosomes through a pathway that largely depends on HrpA. This coupling is not predicted by current models and, thus, inspired us to further investigate the mechanistic link between HrpA and tmRNA.

### Involvement of Hsp15 in ribosome rescue

Recent work has shown that, *in vitro*, HrpA can split stalled 70S ribosomes into 30S and 50S subunits, an activity that depends on the recognition of a ribosome collision interface, attachment to a downstream mRNA segment, and ATP hydrolysis by the HrpA ATPase domain^14^. Such a splitting reaction should, in most cases, produce topologically trapped 50S·peptidyl-tRNA complexes. Although it remains unclear how these are resolved in *E. coli*, a dedicated recognition factor, Hsp15, with high affinity for such complexes is present^26^. Hsp15 is induced during the heat-shock response and stabilizes 50S·peptidyl-tRNA complexes that accumulate under heat stress^25^. Notably, studies have shown that the extent of Hsp15 binding to 50S subunits increases upon chloramphenicol treatment, suggesting that its role is not limited to heat-shock stress but may extend to antibiotic-induced translational stress^26^. We therefore decided to apply single-molecule tracking to characterize the diffusional behavior of Hsp15, to assess its involvement in ribosome rescue pathways, and to test whether its ribosome association is linked to HrpA activity.

To fluorescently label Hsp15, we constructed genomic fusions in which a HaloTag was linked to Hsp15, either at the N terminus or the C terminus, neither of which is predicted to directly contact the 50S subunit based on a high-resolution structural study^25^. Single-molecule tracking was performed using stroboscopic laser illumination (3 ms) at a 5 ms camera exposure. HMM and spatial analysis of single-molecule trajectories revealed the same two principal diffusive states for Hsp15 across all fitted model sizes with more than two states, independent of whether the protein was labelled at the N- or C-terminus (Supplementary Fig. 12, Supplementary Data 31-32). The states included a slow diffusion state with an average diffusion coefficient of approximately 0.3-0.4 µm^2^ s^−1^ and a fast state more than an order of magnitude faster (>5 µm^2^ s^−1^). In models with four or more states, the slow diffusive population could be further subdivided into two distinct substates with diffusion coefficients of approximately 0.06 µm^2^ s^−1^ (slow substate 1), and approximately 0.6 µm^2^ s^−1^ (slow substate 2) (Fig. 3a, Supplementary Fig. 12, Supplementary Data 31-32). Slow substate 1 showed pronounced membrane localization, whereas slow substate 2 displayed a homogeneous distribution throughout the cytoplasm (Fig. 3a). These results are consistent with the expectation that Hsp15 should display diffusion similar to that of free 50S subunits, which diffuse at approximately 0.5 µm^2^ s^−1 28^. However, as a subset of Hsp15 substrates likely corresponds to 50S subunits bound to peptidyl-tRNAs carrying membrane-inserted nascent chains, a membrane-associated fraction is also expected (i.e., slow substrate 1).

**Fig. 3.**
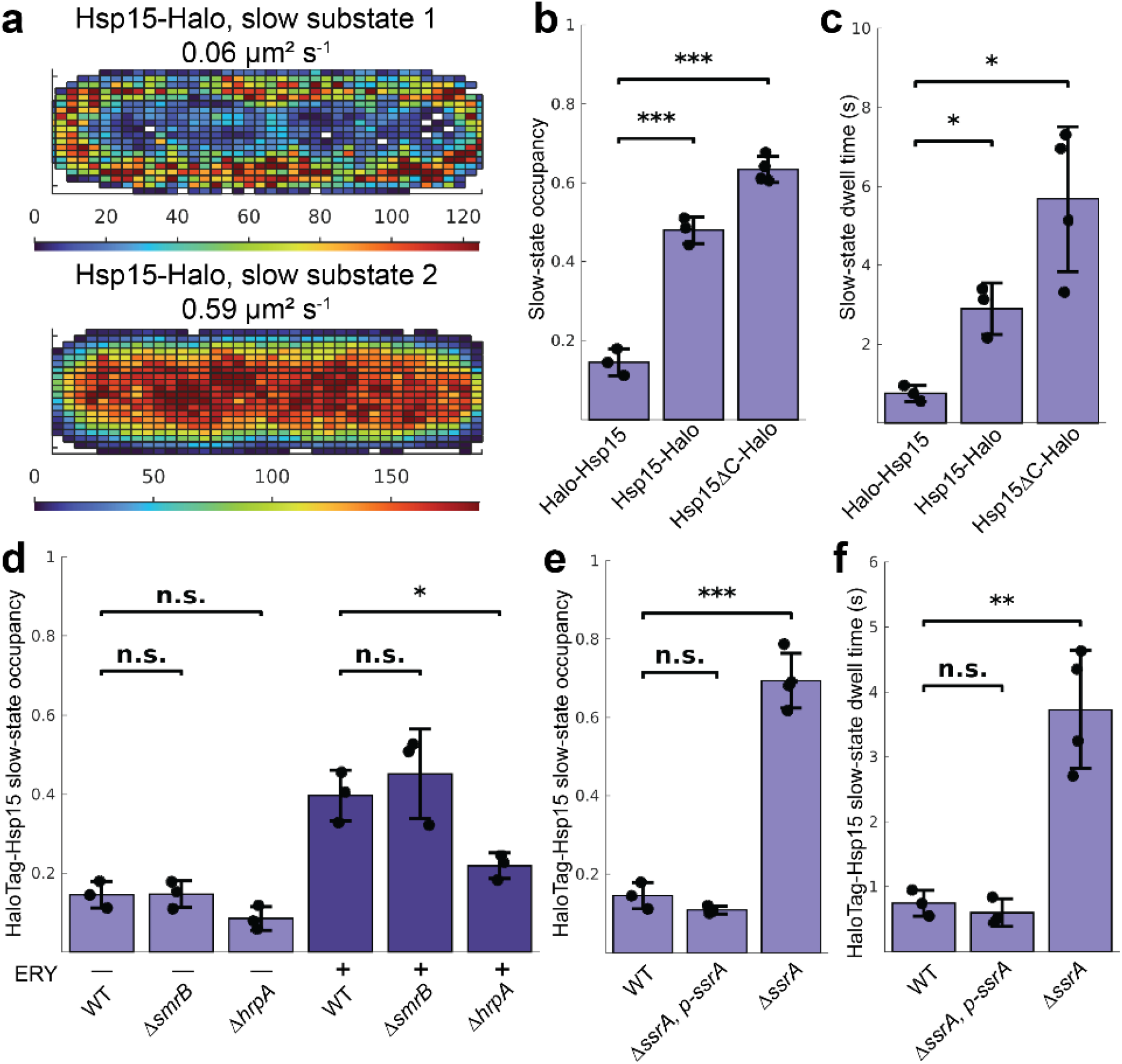
Hsp15 ribosome binding is linked with HrpA and resolved by tmRNA. **a-b**. Ribosome-bound occupancies (**a**) and dwell times (**b**) for N- or C-terminally HaloTag labelled Hsp15, and the effect of removing the C-terminal α-helix in the context of C-terminally labelled Hsp15. **c**. Spatial distributions of Hsp15-HaloTag in slow substates 1 and 2 derived from the 3-state HMM model. The average diffusion coefficients in substates are indicated. **d**. Ribosome-bound state occupancies of HaloTag-Hsp15 in WT, *ΔsmrB* and *ΔhrpA* strains without or with erythromycin treatment. **e-f**. Ribosome-bound occupancies (**e**) and dwell times (**f**) of HaloTag–Hsp15 in WT, *ΔssrA*, and *ΔssrA* complemented with plasmid-encoded *ssrA*. In **b - f**, data are derived from coarse-grained 3-state HMM models. Bars represent averages from independent experiments, and error bars indicate standard deviations between experiments. Statistical significance between strains was assessed using a two-sided unpaired t-test. P-values are indicated in the figure as follows: P < 0.05 (*), P < 0.01 (**), P < 0.001 (***), and not significant (n.s.) otherwise.

Interestingly, Hsp15 labeled via the C-terminus showed significantly longer dwell times in the slow diffusion state and higher steady state occupancy, than the construct labeled via the N-terminus (2.9 ± 0.7 s and 48 ± 3% versus 0.7 ± 0.2 s and 15 ± 3%, respectively) (Fig. 3b-c, Supplementary Data 31-32). Given that under non-stress conditions only a small fraction of Hsp15 is reported to associate with 50S subunits^25^, we considered the possibility that C-terminal labeling interferes with downstream processing of Hsp15-bound ribosomal intermediates. The HaloTag was fused to the C-terminal α-helix of Hsp15, a region of unknown function, via a linker. We tested the role of the C-terminal helix by its removal in the C-terminally labeled Hsp15 construct and then analyzed the effects on the ribosome-binding kinetics. We observed that such a mutation led to further complex stabilization and an additional increase in the slow diffusion state dwell time (5.7 ± 1.8 s) and a steady state occupancy of the slow diffusion state reaching 63 ± 3% (Fig. 3b-c Supplementary Data 31-33). In these experiments, the estimated diffusion coefficients and state occupancies are consistent across different HMM model sizes and labeling strategies for Hsp15 (Supplementary Fig. 13, Supplementary Data 31-33). However, the dwell times show higher variability, particularly, for the Hsp15 mutant lacking the C-terminal α-helix, likely due to the low frequency of state-transition events which prevents accurate dwell time calculations (Supplementary Fig. 13, Supplementary Data 31-33). Overall, these results support the involvement of the Hsp15 C terminus in the processing of aberrant 50S subunits bound to peptidyl-tRNAs, and suggest that steric obstruction or removal of the C-terminal α-helix restricts downstream processing and release of Hsp15.

We next tested whether erythromycin treatment leads to an increased production of substrates for Hsp15 binding. Based on our observation that C-terminal labeling impairs efficient downstream processing of Hsp15-bound ribosomes, all subsequent experiments were performed using the N-terminal HaloTag fusion of Hsp15. Upon erythromycin treatment, we observed a pronounced increase in the combined slow diffusion states corresponding to ribosome-bound Hsp15, from 15 ± 3% to 40 ± 6% (Fig. 3d, Supplementary Data 31,34). In line with previous reports showing an increased association of Hsp15 with the 50S subunit under chloramphenicol treatment, these results suggest that, in addition to its role in heat-shock response, Hsp15 binds aberrant ribosomes generated during antibiotic treatment.

To assess how Hsp15 engagement with aberrant ribosomes depends on ribosome rescue pathways, we examined Hsp15 ribosome binding dynamics in *ΔsmrB* and *ΔhrpA* strains. This allowed us to test whether the activity of Hsp15 is functionally linked to the collision-sensing factor HrpA, which we expect to convert collided ribosomes into substrates for Hsp15. We found that, in the *ΔhrpA* strain, the fraction of Hsp15 bound to ribosomes was reduced compared to in the WT, decreasing modestly from 15 ± 3% to 9 ± 3% (P = 0.084) under untreated conditions, and more markedly in the presence of erythromycin from 40 ± 6% to 22 ± 6% (P = 0.024) (Fig. 3d, Supplementary Data 31,34-36). No corresponding decrease was observed in the *ΔsmrB* strain (Fig. 3d, Supplementary Data 31,34,37-38). Notably, the cellular abundance of Hsp15 was unchanged in strains lacking *smrB* or *hrpA* compared to WT and was unaffected by erythromycin treatment (Supplementary Fig. 14).

In summary, single-molecule tracking of Hsp15 indicates that erythromycin treatment promotes the formation of aberrant ribosomes that serve as substrates for Hsp15 binding, and that the generation of these substrates is linked to the ribosome collision factor HrpA.

### tmRNA processes Hsp15-bound ribosomes

The experiments presented above showed that, under erythromycin treatment, HrpA generates substrates for both tmRNA and Hsp15, suggesting that both factors act downstream of ribosome collisions processed by HrpA. This raises the possibility that Hsp15 and tmRNA are functionally linked. We therefore tested how deletion of tmRNA affects the ribosome-binding kinetics of Hsp15 and, conversely, how deletion of Hsp15 influences the ribosome-binding kinetics of tmRNA.

While the deletion of Hsp15 had no strong effect on the ribosome-binding kinetics of tmRNA (Supplementary Fig. 15, Supplementary Data 19,22, 39-40), strikingly, deletion of the *ssrA* gene (i.e., tmRNA) caused HaloTag–Hsp15 to predominantly occupy ribosome-bound states even in the absence of antibiotic treatment, with the bound fraction increasing from 15 ± 3% in WT background to 69 ± 7% in the *ΔssrA* strain (Fig. 3e, Supplementary Data 31, 41). The dwell time in the ribosome-bound state showed a corresponding increase, from 0.7 ± 0.2 s to 3.7 ± 0.9 s (Fig. 3f, Supplementary Data 31, 41). This accumulation indicates that, in the absence of tmRNA, Hsp15-bound ribosomes cannot be efficiently processed, and Hsp15 remains ribosome-bound. Reintroduction of *ssrA* on a low-copy-number plasmid in *ΔssrA* cells reduced the occupancy and dwell time of ribosome-bound Hsp15 to levels slightly below those observed in wild-type cells (Fig. 3e-f, Supplementary Data 31, 42).

Previous studies have proposed that processing of Hsp15-bound ribosomes may be mediated by ArfB, PrfH, or Pth, that would cleave the bond between the nascent peptide and the stuck P-site tRNA ^43,44^. Given that *pth* is essential for bacterial growth and hence cannot be deleted and that *prfH* already carries a large inactivating deletion in *E. coli* laboratory strains, we instead tested whether overexpression of these two proteins as well as of ArfB accelerates processing of Hsp15-bound ribosomes (Supplementary Fig. 16). Single-molecule tracking of HaloTag-Hsp15 in a *ΔssrA* strain showed that only overexpression of ArfB caused a moderate decrease in ribosome-bound occupancy of HaloTag-Hsp15 but without significant effect on the dwell-time (Supplementary Fig. 16a-b, Supplementary Data 41, 43). However, deletion of *arfB in ssrA*^*+*^ *cells* had no detectable effect on Hsp15 ribosome-binding kinetics (Supplementary Fig. 16c, Supplementary Data 31, 44). Overexpression of Pth led to a modest reduction in dwell time without affecting occupancy, whereas PrfH overexpression affected neither dwell time nor occupancy (Supplementary Fig. 16a-b, Supplementary Data 41, 45-46). These effects were small compared to those observed upon tmRNA expression, suggesting that ArfB, Pth and PrfH are unlikely to play a physiological role in the processing of Hsp15-bound ribosomes.

These results showed that Hsp15-bound ribosomes are processed predominantly by tmRNA. Since tmRNA can only target 70S ribosomes, Hsp15-bound ribosomes are expected to pass through a 70S-associated intermediate accessible to both Hsp15 and tmRNA. Although Hsp15 does not associate with actively translating 70S ribosomes^26^, analysis of available structures of 50S·peptidyl-tRNA·Hsp15 complexes and 70S ribosomes, together with AlphaFold modelling, suggested that Hsp15 binding is compatible with a non-translating 70S context lacking mRNA in the E-site channel, with the C-terminal α-helix of Hsp15 occupying this channel, positioned at the interface between 16S rRNA, uS7, and uS11 (Supplementary Fig. 17). To test whether such contacts occur in cells, we expressed a variant of Hsp15 carrying the unnatural photo-crosslinkable amino acid p-benzoyl-L-phenylalanine (BPA) at position 124 within the C-terminal α-helix. Upon UV irradiation of cells expressing such a construct, we observed crosslink products with uS7, supporting that Hsp15 contacts not only the 50S subunit but also the 30S subunit (Supplementary Fig. 18).

### Cryo-EM analysis of Hsp15-bound ribosomes isolated from cells under erythromycin treatment

Given that in the *ΔssrA* strain we observed a pronounced enrichment of Hsp15 in the slow diffusive state, which is expected to contain 70S complexes according to our prediction, we performed a pull-down using N-terminally His-tagged Hsp15, concentrated the ribosomes through pelleting and analyzed the recovered material by cryo-EM.

Image processing identified 1.8 M ribosomal particles, with a roughly equal number of 50S and 70S particles after global 3D classification (Supplementary Fig. 19). The most populated 50S class (514,726 particles, 2.23 Å) consisted of a 50S·peptidyl-tRNA·Hsp15 complex with a peptidyl-tRNA bound at the 50S P-site and Hsp15 interacting with the tRNA anticodon stem and 23S helices H68 and H69 (Fig. 4a-b; Supplementary Figs. 20, 21). The majority of the 70S particles contained a peptidyl-tRNA and had an unrotated 30S subunit. Focused 3D classification with a mask covering the P-site tRNA and the N-terminal domain of Hsp15 followed by a second classification with a mask also including the 30S head, identified several 70S complexes with Hsp15 (Supplementary Fig. 19). Notably, density for a nascent chain attached to the P-site tRNA was visible throughout the exit tunnel in all the Hsp15-bound 50S and 70S complexes (Fig. 4a).

**Fig. 4.**
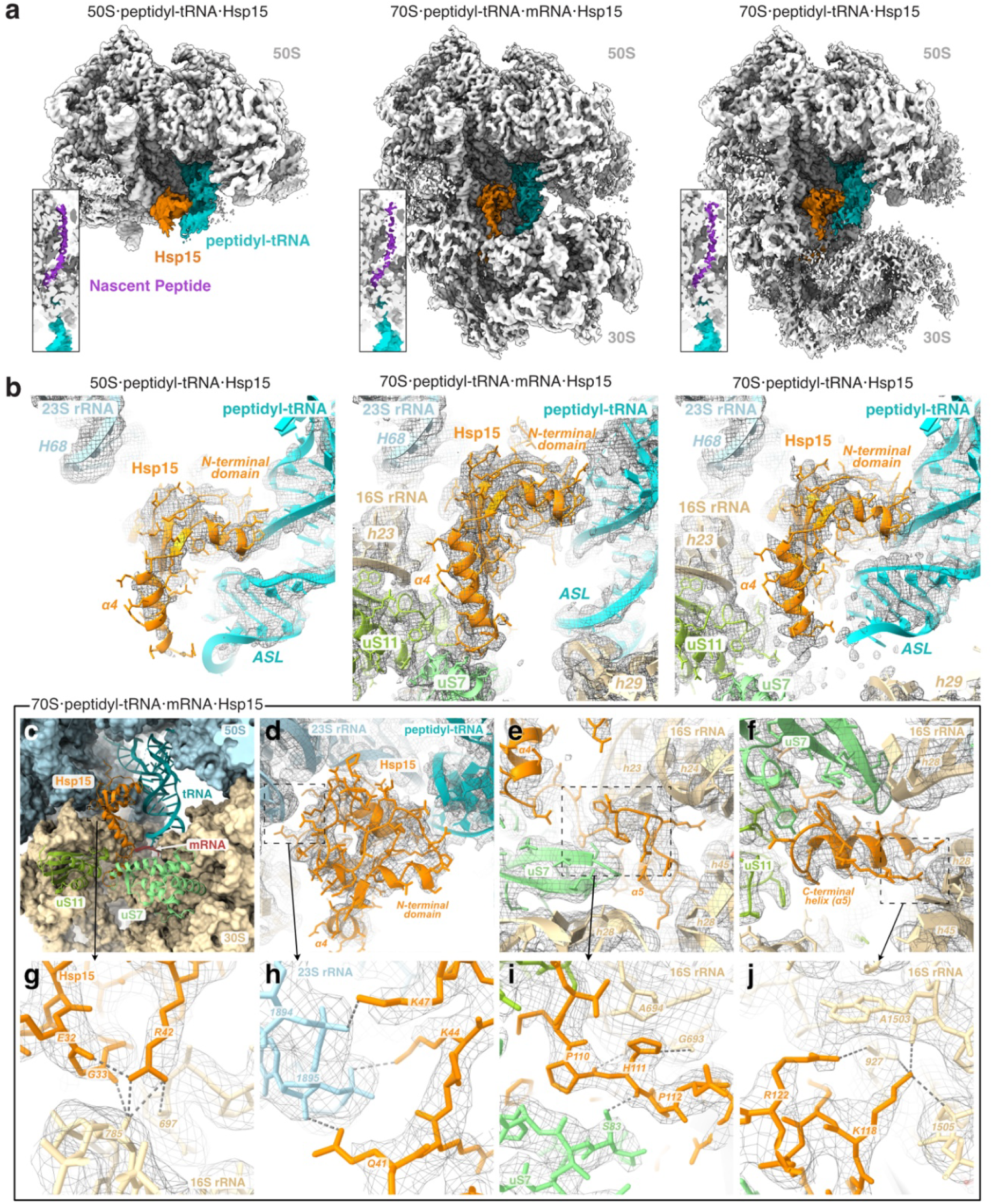
Cryo-EM reconstruction of ribosomal Hsp15 complexes. **a**. left, 50S complex with peptidyl-tRNA (turquoise) and Hsp15 (orange) with inset showing density in the exit tunnel for the nascent peptide (purple), middle, 70S complex with the 30S and P-site tRNA in classical conformation and mRNA at the P-site, right, 70S complex with the small subunit head disengaged from the central protuberance of the 50S and the peptidyl-tRNA anticodon stem not bound to the 30S. **b**. Close-ups of the N-terminal domain of Hsp15 and anti-codon stem loop of peptidyl-tRNA with cryo-EM map shown as mesh. **c-j**. Details of 70S·peptidyl-tRNA·mRNA·Hsp15 complex. **c**. Hsp15 binding-site relative to the mRNA (red), uS7 (mint green) and uS11 (yellow green). **d**. N-terminal domain. **e**. α4–α5 loop. **f**. C-terminal helix. **g**. N-terminal domain contacts to 16S rRNA (wheat color). **h**. N-terminal domain contacts to 23S rRNA (light blue). **i**. α4–α5 loop contacts to uS7 and 16S rRNA. **j**. C-terminal helix contacts to 16S.

The 50S·peptidyl-tRNA·Hsp15 complex is equivalent to the structure previously determined by cryo-EM^44^. Hsp15 stabilizes the tRNA in a P-site-like state with the anticodon stem loop (ASL) shifted towards the E site (Fig. 4b). Only the N-terminal domain (NTD) and α4-helix of Hsp15 (amino acids 8-102) are resolved, while the C-terminus is flexible (Supplementary Fig. 21a,b). Positively charged residues in the NTD (Arg10, Lys13, Arg24, Arg28, and Lys44) form contacts with the RNA backbone of helices H69 and H68 in 23S rRNA and to the tRNA ASL (Lys22, Arg76 and Arg19) (Supplementary Fig. 21c). The α4-helix is engaged with the anti-codon loop (33-36) of the peptidyl-tRNA (Fig. 4b). Surprisingly, the 50S·peptidyl-tRNA·Hsp15 complex shows an additional well-resolved density near ribosomal protein uL14, which we could assign to the ribosome silencing factor S (RsfS) (Supplementary Fig. 22). RsfS binds a well-documented site near uL14, bL19 and 23S helix H95^45–47^ and inhibits subunit association by sterically hindering interaction of 50S with 16S helix h14 to form the inter-subunit bridge B8.

From the final round of classification, we identified four 70S·Hsp15·peptidyl-tRNA classes: one lacking mRNA and three containing mRNA base-paired to peptidyl-tRNA. Among the mRNA-containing classes, one (41,043 particles, 2.78 Å, Supplementary Fig. 19) provided map quality which enabled unambiguous modelling of all but the first five residues of Hsp15 (Fig. 4d-f). We refer to this complex as 70S·peptidyl-tRNA·mRNA·Hsp15. The other two classes showed weaker density for Hsp15 but exhibited a similar overall architecture (Supplementary Fig. 23).

The 70S·peptidyl-tRNA·mRNA·Hsp15 complex shows Hsp15 bound in a similar location as in the 50S complex, but the α4-helix is disengaged from the anti-codon loop of the peptidyl-tRNA and instead contacts uS7 and uS11 of 30S (Fig. 4e,i), and the NTD binds further up on the ASL (Fig. 4b). The tRNA is in classical P/P state, with the ASL bound to the P-site of the unrotated 30S (Fig. 4a). There is clear mRNA density at the P-site (Supplementary Fig. 24a), but not upstream or downstream. Interestingly, the codon-anticodon helix shows a clear pattern of purines and pyrimidines, suggesting that the ribosomal sample is enriched for certain tRNAs (Supplementary Fig. 24b). The purine-pyrimidine pattern of the codon-anticodon base pairs would be consistent with Gly, Ala, Ser or Asp tRNAs. Absence of a long variable loop, characteristic of tRNA^Ser^, and the purine-pyrimidine pattern in the anticodon stem (Supplementary Fig. 24c) prompted us to model the peptidyl-tRNA as tRNA^Asp^ in cognate interaction with a GAC codon. The NTD of Hsp15 is involved in contacts with both the 23S and the 16S rRNA (Fig. 4g,h). Arg42 makes polar contacts with 16S nucleotides U697 and G785, while the carbonyl oxygens of Glu32 and Gly33 help stabilize the conformation of the Arg42 sidechain (Fig. 4c,g), contrary to the prediction that Arg42 and Glu32 would sterically clash with the 30S subunit^44^.

The C-terminus of Hsp15, which is disordered in the 50S complex, is folded into an α-helix (α5) and occupies the mRNA channel of the 30S subunit in the 70S·peptidyl-tRNA·mRNA·Hsp15 complex, where it blocks the E site (Supplementary Fig. 25). The α4–α5 loop that connects the NTD to the α5-helix, enters the 30S using a Pro-His-Pro motif, forming hydrogen bonds with Ser83 of uS7 and G693 of 16S rRNA (Fig. 4e,i). The C-terminal α5-helix forms contacts with the 16S rRNA helices h28 and h45 in the 30S head as well as with uS7 and uS11 (Fig. 4c,f), in agreement with our observed crosslinking to uS7. Inside the mRNA channel, positively charged residues Arg115, Lys118, Lys119, Arg122 and Arg126 stabilize binding in the negatively charged environment (Supplementary Fig. 25). The guanidium moiety of Arg122 stacks with the A1503 base of 16S rRNA, and Lys118 forms a hydrogen bond to the 2’ hydroxyl group of A1503 (Fig. 4f,j).

Finally, we find a 70S·peptidyl-tRNA·Hsp15 complex which lacks mRNA, but shows density for the C-terminal α5-helix of Hsp15 positioned in the mRNA channel in the same way as in the 70S·peptidyl-tRNA·mRNA·Hsp15 complex (Supplementary Fig. 26b). The peptidyl-tRNA and Hsp15, however, are in conformations closely resembling the 50S·peptidyl-tRNA·Hsp15 complex (Fig. 4a,b). In this complex, the 30S head is unlocked from the 50S resulting in higher mobility and lower local resolution (Supplementary Fig. 20c). The α4–α5 loop of Hsp15 is disordered, as well as the β-hairpin (73–90) of uS7 and the anticodon loop of the peptidyl-tRNA (Supplementary Figs. 21a, 26a). Comparison of all complexes shows similar Hsp15 conformations apart from the flexible C-terminus (Supplementary Fig. 21b).

## DISCUSSION

The large ribosomal subunit carrying a trapped peptidyl-tRNA is a key intermediate in ribosome rescue across all domains of life. In eukaryotes, these are processed by the RQC pathway. While some bacteria encode an RqcH–RqcP system analogous to the eukaryotic RQC, many, including *E. coli*, lack RqcH and instead encode only an RqcP homolog, which can bind but not by itself process 50S·peptidyl-tRNA complexes. In *E. coli*, the RqcP homolog, Hsp15, is strongly upregulated during heat stress, when 50S·peptidyl-tRNA accumulates. Ribosome stalling with subsequent ribosome collisions represent another source of these aberrant complexes, where the recently identified collision factor HrpA was shown to convert stalled ribosomes into 50S·peptidyl-tRNA *in vitro*. Thus, understanding how 50S·peptidyl-tRNA is resolved in the absence of RqcH is important for understanding ribosome rescue in *E. coli* and other bacteria.

Based on the results presented, we propose a model linking tmRNA to the processing of 50S·peptidyl-tRNA complexes generated by HrpA during ribosome collisions and likely from other sources, such as heat stress (Fig. 5). Consistent with this model, we find that Hsp15-bound ribosome species are predominantly processed by tmRNA (Fig. 3), indicating that they are directed into the trans-translation pathway. Since tmRNA requires the full 70S ribosome for processing, our results implied a 70S·peptidyl-tRNA·Hsp15 intermediate. Crosslinking and cryo-EM analysis of Hsp15-bound ribosomes from tmRNA-deficient cells with erythromycin-induced stalling shows that Hsp15 binds not only 50S·peptidyl-tRNA, as previously believed^26,43,44^, but also a range of 70S·peptidyl-tRNA complexes. In these complexes, the C-terminal α-helix of Hsp15 occupies the free E-site mRNA channel (Fig. 4f), explaining why Hsp15 does not associate with actively translating 70S ribosomes. We captured 70S·peptidyl-tRNA·Hsp15 complexes lacking mRNA, as well as complexes with mRNA visible in the P site, but either truncated or displaced from its canonical path, where the E site is occupied by the Hsp15 C-terminus. Given that an empty A-site mRNA channel is the key determinant for tmRNA recognition, 70S·peptidyl-tRNA·Hsp15 complexes lacking mRNA constitute accessible substrates for tmRNA in ssrA^+^ cells. 70S·peptidyl-tRNA·Hsp15·mRNA complexes with truncated or displaced mRNA may likewise be accessible to tmRNA. We suggest that all of these 70S complexes could arise either from full or partial displacement of mRNA from translating ribosomes or from reassociation of free 30S subunits with 50S·peptidyl-tRNA. The latter would also explain how dissociated 30S and 50S·peptidyl-tRNA, for example during heat shock, can enter the trans-translation pathway. In this case, the C-terminal α-helix of Hsp15 could facilitate binding of mRNA-free 30S subunits to 50S·peptidyl-tRNA by providing an additional contact and thereby stabilizing 70S·peptidyl-tRNA complexes lacking mRNA.

**Fig. 5.**
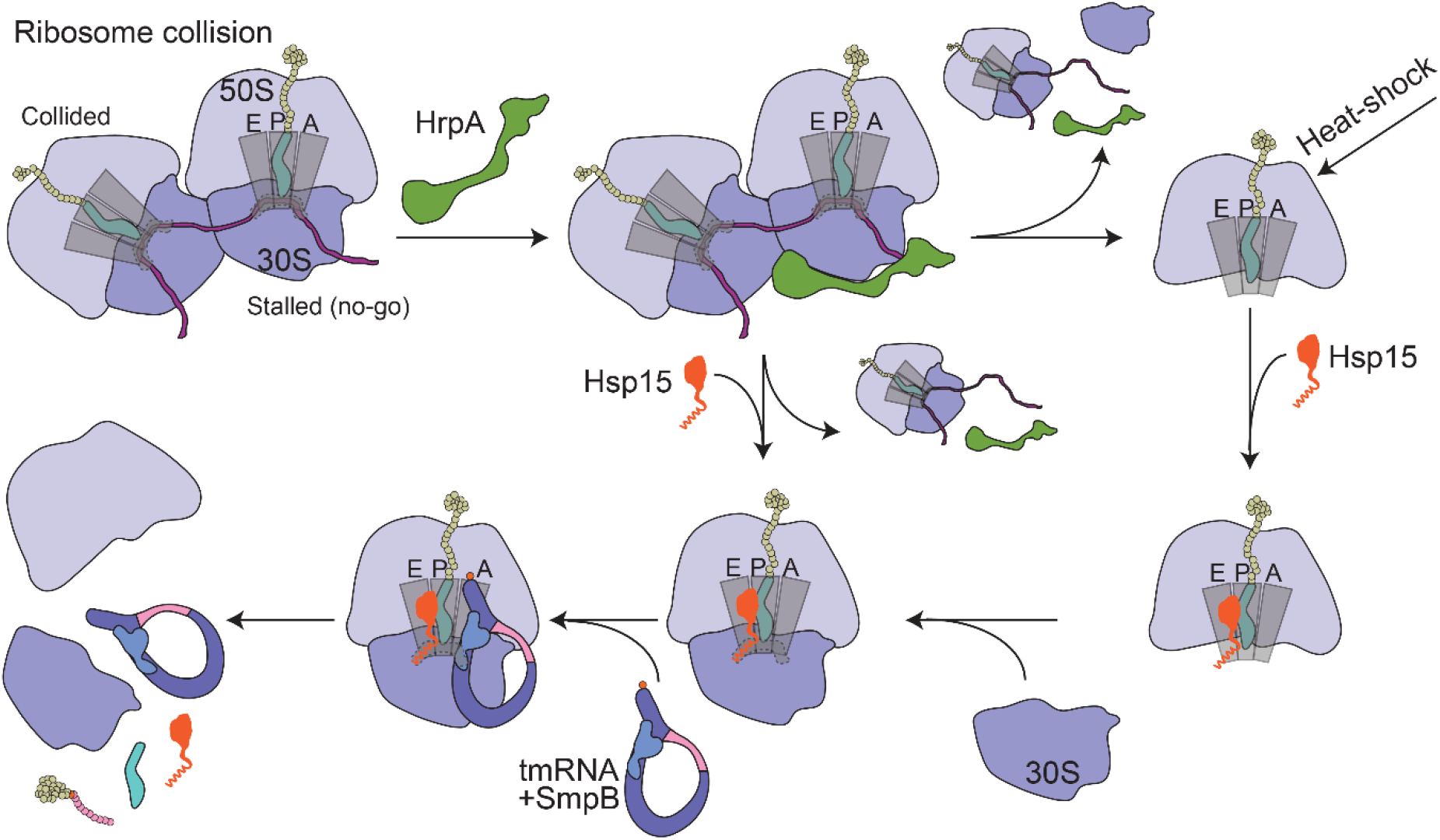
Proposed model for tmRNA- and Hsp15-mediated rescue of stalled ribosomes. Ribosome collisions are recognized by HrpA, which processes stalled 70S ribosomes either by subunit splitting (supported by ^14^) or by displacement of mRNA. This generates 50S·peptidyl-tRNA intermediates, which may also arise under heat stress, or 70S·peptidyl-tRNA complexes. Hsp15 binds these species, with its extended C-terminal α-helix serving as a recognition element for 70S·peptidyl-tRNA complexes lacking mRNA in the E-site channel and potentially promoting reassociation with mRNA-free 30S subunits. The resulting 70S·peptidyl-tRNA·Hsp15 complexes are then directed to the tmRNA-mediated pathway, leading to ribosome rescue and recycling.

These results, in addition, provide a mechanistic basis for our first key finding that, under erythromycin treatment, the collision factor HrpA functions upstream of tmRNA, generating intermediates for tmRNA-mediated rescue (Fig. 2c). While the mechanistic details of HrpA activity remain to be fully resolved, available data^14^ are in line with our results: *In vitro*, in the presence of excess IF3 which prevents subunit reassociation, HrpA splits stalled 70S ribosomes into subunits, generating 50S·peptidyl-tRNA complexes. *In vivo*, however, HrpA action leads to accumulation of 70S monosomes, a result suggesting either reassociation of 30S to the 50S·peptidyl-tRNA complex, or that HrpA *in vivo* merely displaces the mRNA on a 70S complex, rather than splitting the subunits. Both scenarios are compatible with our results, and importantly, suggest that these intermediates can be directed to the trans-translation pathway.

Thus, from our single-molecule tracking studies of ribosome rescue factors, backed up by cross-linking and cryo-EM structures, we have found that aberrant stalled ribosomes can enter the general tmRNA pathway through a mechanism involving the recently found collision factor HrpA and the RqcP analogue Hsp15. Interestingly, although our findings point to an even broader role for tmRNA as a universal ribosome rescuer, bacteria have nevertheless evolved multiple redundant rescue mechanisms, including alternative rescue factors and RQC‐like pathways. This redundancy highlights the critical importance of efficient ribosome rescue and reflects the severe threat that stalled ribosomes pose to the cell.

In addition to this key finding, our results suggest that all rescue factors investigated, i.e., tmRNA, SmrB, and HrpA, are continuously binding and processing stalled ribosomes also under normal growth conditions (Fig. 2a). Thus, ribosome stalling is probably an unavoidable general feature of protein synthesis. Upon treatment with a sub-lethal dose of erythromycin, however, the usage frequency of all factors is drastically elevated.

Although a link between SmrB and tmRNA has been reported previously using reporter constructs with stalling sequences, we do not detect a significant effect of SmrB deletion on global tmRNA recruitment under erythromycin treatment (Fig. 2c). SmrB has been shown to act preferentially on higher-order collisions involving more than two ribosomes^16^, whereas under our conditions of moderate antibiotic stress, HrpA likely resolves the majority of collision events. In addition, SmrB cleaves mRNA and promotes its decay, which may reduce the formation of substrates that would otherwise be processed by tmRNA. Furthermore, SmrB does not have a strong effect on antibiotic sensitivity, suggesting that it might have a different or more specialized role, for example, in gene regulation.

## METHODS

### Bacterial strains and growth conditions

All constructed strains were derived from *Escherichia coli* K-12, strain MG1655, which is referred to as the wild-type (WT) strain. The *prfH* gene was amplified from genomic DNA of *E. coli* strain DA33135, a gift from Dan I. Andersson^48^.

### Plasmid construction

For all molecular cloning procedures, Q5 High-Fidelity DNA Polymerase (NEB) was used for PCR amplification following the manufacturer’s protocols. PCR products were treated with FastDigest DpnI (Thermo Scientific) and purified using the E.Z.N.A. Cycle Pure Kit (Omega BIO-TEK) following the manufacturer’s protocols. Gibson assembly was performed according to the manufacturer’s instructions using NEBuilder HiFi DNA Assembly Master Mix (NEB). Ligations were performed using T4 DNA ligase (NEB).

Two strategies were used for plasmid construction: (i) Gibson assembly of DNA fragments obtained by PCR, and (ii) PCR amplification of the parental plasmid using mutagenic oligonucleotides, followed by re-circularization of the linear PCR product through end phosphorylation with T4 polynucleotide kinase (NEB) and ligation using T4 DNA ligase. The oligonucleotides used in this study are listed in Supplementary Data 47. Plasmids used in this study and cloning procedures for plasmids constructed in this study are provided in Supplementary Data 48.

### Construction of strains carrying chromosomal mutations

Chromosomal mutations, including deletions, fusions, and gene replacements, were generated using λ Red–mediated recombineering by electroporation of linear PCR-amplified DNA fragments, or single-stranded oligonucleotides, followed by selection on LA plates under appropriate selective conditions (antibiotics or sucrose). The λ Red genes were expressed from plasmid pSIM5-TET^49^ upon induction by a temperature shift to 42 °C. The oligonucleotides and template DNAs used for PCR amplification are listed in Supplementary Data 47. All strains generated in this study, together with descriptions of the cloning procedures, are provided in Supplementary Data 49.

A previously described scar-free genome editing strategy was used to (i) replace the chromosomal *ssrA* gene with a variant carrying the MS2 aptamer inserted at position 175, including versions carrying the G3A or A86C mutations; (ii) delete *smpB* gene in tmRNA-MS2^g^ strain (iii) introduce an N-terminal HaloTag fusion to Hsp15 encoded by hslR; and (iv) generate a C-terminal HaloTag fusion to rpsG^50,51^.

### Doubling time measurements

Doubling time measurements were performed to assess the functionality of the MS2-tagged tmRNA and the impact of tmRNA mutations on growth. Strains were streaked on LB agar plates without antibiotics (WT, tmRNA-MS2^g^, *ΔsmpB* tmRNA-MS2^g^, tmRNA-G3A-MS2^g^, and tmRNA-A86C-MS2^g^) or on plates containing 50 µg ml^−1^ kanamycin (*ΔssrA::kan* or *ΔsmpB::kan*) and incubated overnight at 37 °C. Three independent colonies per strain were used to inoculate 3 ml LB with or without kanamycin (50 µg ml^−1^), as appropriate, and grown to approximately OD_600_ ≈ 1. Cultures were diluted 1:10000 in LB with or without erythromycin (10 or 25 µg ml^−1^). Aliquots (150 µl) were transferred to 96-well plates, and OD_600_ was recorded every 5 min using a BioTek Synergy H1 plate reader with continuous shaking. Doubling times were determined using a custom MATLAB script^50^ which fits an exponential growth function to the exponential part of the growth curve. The experiment was repeated four times and the doubling time per strain was calculated averaging the results from all the twelve colonies and the standard deviation was calculated.

### *In vivo* labeling of HaloTag fusion proteins

*In vivo* fluorescence labeling of HaloTag fusion proteins was performed as described previously^29^, with minor modifications. Cell cultures used for labeling were prepared either by diluting overnight cultures 1:100 into fresh LB (15 ml) supplemented with the appropriate antibiotics and growing to an OD_600_ of 0.6–1.0, or by inoculating fresh LB (15 ml) with colonies from overnight LB agar plates and growing to the same optical density. The cells were harvested by centrifugation at 4000 g for 5 min and washed twice with 1 ml M9 medium with 0.4% glucose. Labeling was performed at 25 °C for 30 min in 150 µl EZ Rich Defined Medium (RDM; Teknova) supplemented with JFX549 HaloTag ligand (a gift from the Lavis lab) at final concentrations of 0.2 µM for L9–HaloTag, 1 µM for HaloTag fusions with Hsp15, and 3.1 µM for labeling of tmRNA, HaloTag–HrpA, and HaloTag-SmrB. After labeling, cells were washed four times in 1 ml M9 medium supplemented with 0.4% glucose, incubated for 30 min at 37 °C in 3 ml RDM with shaking, and subsequently washed three additional times in M9 medium containing 0.4% glucose. Labelled cells were resuspended in 200 µl of M9 medium containing 0.4% glucose and either used immediately for microscopy or stored at −80 °C in 10% glycerol. For microscopy experiments cell aliquots were resuspended to OD600 of ≈0.003 in M9 medium with 0.4% glucose and applied onto a 2% agarose pad (SeaPlaque GTG Agarose, Lonza) in RDM medium. The agarose pad was cast within a Gene Frame (Thermo Fisher) and sealed between a microscope slide and a #1.5H coverslip (Thorlabs). The sample was placed on the microscope within an incubation chamber maintained at 37 ± 2 °C. For experiments involving erythromycin treatment, the antibiotic was added to the agarose solution used to prepare the imaging pad to a final concentration of 25 µg ml^−1^. For experiments involving expression of *arfB, pth*, or *prfH* from pQE plasmids, IPTG was added to the agarose pads at a final concentration of 50 µM. Cells were allowed to grow for approximately 70–120 min to form mini-colonies, after which image acquisition was performed.

All sample preparation and subsequent microscopy experiments were independently repeated at least three times, with the exception of L9–HaloTag experiments, which were performed twice.

### Optical setup

Widefield epifluorescence microscopy was performed using an inverted microscope (Nikon Ti2-E) equipped with a CFI Plan Apo λ 100×/1.45 NA objective (Nikon). The microscope was enclosed in an incubation chamber (H201-ENCLOSURE, OKOlab) equipped with a temperature controller (H201-T-UNIT-BL, OKOlab), maintaining the temperature at 37 ± 2 °C. Phase-contrast and bright-field images, as well as fluorescence time-lapse movies, were acquired using an Orca Quest camera (Hamamatsu). Fluorescent molecules were excited using a 546 nm laser (2RU-VFL-P-2000-546-B1R, 2000 mW; MPB Communications) at a power density of 3 kW cm^−2^ at the sample plane. Illumination was applied in stroboscopic mode with one 3 ms laser pulse per 30 ms camera exposure for tmRNA-MS2^g^, tmRNA-MS2^p^, HaloTag–HrpA, and HaloTag-SmrB tracking, and with one 3 ms laser pulse per 5 ms camera exposure for Hsp15 HaloTag fusions. For each mini-colony, 300–700 fluorescence images were recorded to generate time-lapse movies. Microscope operation was controlled via μManager, and image acquisition from multiple positions was carried out using custom μManager plugins.

### Analysis of microscopy data

Image analysis was performed using a custom MATLAB-based pipeline, as described previously^28,29,32^. Briefly, phase-contrast images were used to generate segmentation masks of individual cells using the Per-Object Ellipse Fit (POE) method for adaptive thresholding^52^. Segmentation masks were minimally curated manually to remove poorly segmented cells. Single fluorophores were detected in fluorescence images using a radial symmetry-based detection algorithm^53^. Dot localizations were refined using symmetric Gaussian point spread function (PSF) modelling and maximum a posteriori fitting, which also provided estimates of localization uncertainty that were used to exclude unreliable dots^54^. Two-dimensional diffusion trajectories of individual molecules were constructed using the uTrack algorithm^55^. Trajectory reconstruction was restricted to segmented cells. Trajectory building within segmented cells was initiated only when ≤ 1 fluorophore was detected per cell in consecutive frames.

The resulting trajectories were analyzed using a previously described Hidden Markov model (HMM)-based approach^32,54^. The algorithm explicitly incorporates both spot coordinates and their associated localization uncertainties and accounts for missing data points. For each experiment, all trajectories longer than five frames were collectively fitted to a multi-state diffusion model with a predefined number of states, each defined by its diffusion coefficient, occupancy, and dwell time. All datasets were fitted to model sizes ranging from 2 to 8 diffusion states. Larger models were coarse-grained to 2-state representations based on clustering of diffusion states using a threshold at 0.7 µm^2^ s^−1^ (for tmRNA-MS2^g^, tmRNA-MS2^p^, HaloTag-HrpA, and HaloTag-SmrB), or 1 µm^2^ s^−1^ (for Hsp15 and its mutants fused to HaloTag) to separate “Slow diffusion state” and “Fast diffusion state”. For Hsp15, coarse-graining was additionally performed using a three-state scheme with thresholds at 0.25 and 1 µm^2^ s^−1^, resolving two distinct slow diffusive substates. Occupancies and dwell-times presented in bar plots were calculated from 3-state HMM models coarse-grained to two states, with averages calculated from independent experiments. Error bars represent standard deviations between these independent experiments. Statistical significance between groups was assessed using a two-sided unpaired t-test. P-values are indicated in the figures as follows: P<0.05 (*), P<0.01 (**), P<0.001 (***), and not significant (n.s.) otherwise.

Spatial distributions were calculated from three-state HMM models following coarse-graining into two states and displayed as heat maps using normalized cell coordinates.

### Quantification of tmRNA by northern blotting

Strains in which tmRNA abundance was to be quantified were inoculated into 5 ml LB and incubated overnight at 37 °C. An aliquot (75 µl) of each overnight culture was diluted into 15 ml LB, with or without erythromycin at 25 µg ml^−1^ and incubated at 37 °C until reaching an OD_600_ of 0.4–0.6. Cells were collected from 2 ml of culture by centrifugation at 10, 000 × g. for 1 min. Cell pellets were either stored at −20 °C or immediately processed for RNA extraction according to a previously described protocol^56^. Briefly, cell pellets were resuspended in 320 µl RNASwift lysis reagent (4% SDS, 0.5 M NaCl,) prewarmed to 37 ◦C. Samples were incubated at 37 °C for at least 3 min with occasional mixing by tube inversion, followed by addition of 160 µl 5 M NaCl. Precipitates were removed by two rounds of centrifugation at 16,000 × g for 4 min. An aliquot (300 µl) of the resulting supernatant was mixed with 300 µl ice-cold isopropanol and incubated overnight at −20 °C to precipitate RNA. The RNA was pelleted by centrifugation (18,000 × g., 20 min, 4 °C), washed with 70% ice-cold ethanol, air-dried, and resuspended in 50 µl Milli-Q water. RNA concentrations were determined (1 A_260_ ≈ 40 ng µl^−1^), samples were adjusted to 80 ng µl^−1^, mixed 1:1 with 2× TBE-urea buffer (BIO-RAD), and heated at 70 °C for 5 min. From each sample 10 µl (∼400 ng of RNA) was loaded onto two 5% polyacrylamide TBE-Urea gels (19:1 acrylamide/bis-acrylamide, 7 M urea) and separated for 35 min at 200 V. The two gels were each blotted at 30 V for 1 hour using 0.5xTBE buffer onto an Amersham™ Hybond-N+ nylon membrane (Cytiva). Membranes were fixed by incubation for 2 hours at 80 ◦C before blocking by incubation in 15 ml Recette-Church buffer (0.25 M sodium phosphate buffer pH 7.2, 1 mM EDTA, 7% (w/v) SDS, 0.5% (w/v) BSA, 0.004 mg/ml salmon sperm DNA) for 2 hours at 37 ◦C in a tube rotator. IR-dye labeled DNA oligos specific for tmRNA (/5IRD700/cggaggctagggagagagggc) and 5S rRNA (/5IRD800/ggcgctacggcgtttcacttc) were added (1 nmol each) and the incubation was continued overnight at 37 ◦C. Membranes were washed at 37 ◦C twice with 20 ml 2xSSC buffer with 2% SDS and then once with 40 ml 1xSSC buffer with 1% SDS, each wash for about 30 minutes. Bands corresponding to tmRNA and 5S rRNA were detected using a Licor Odyssey CLx Imager and quantified using the Image Studio software. The tmRNA signal of each sample was normalized by using the corresponding 5S signal as an internal standard, equalizing the 5S signal for all samples from four experiments and weighing the tmRNA signal accordingly. The tmRNA signal was averaged over the two gels from each experiment and the mean of the four resulting values per strain or condition and the standard deviation were calculated. The relative abundancies of tmRNA in all samples with respect to the mean abundance in tmRNA-MS2^g^ were calculated and plotted along with the four individual measurements.

### Quantification of relative HaloTag-Hsp15 abundance

Strains *E. coli HaloTag–Hsp15, E. coli ΔhrpA::kan HaloTag–Hsp15, and E. coli ΔsmrB::kan HaloTag–Hsp15* used for quantification of HaloTag–Hsp15 were grown overnight in 5 ml LB at 37 °C. An aliquot (100 µl) of each overnight culture was diluted into 10 ml LB and incubated at 37 °C until reaching an OD_600_ of approximately 0.5. An aliquot (0.75 ml) of each culture was diluted into 15 ml LB with or without erythromycin at 25 µg ml^−1^ and incubated at 37 °C until reaching an OD_600_ of 0.5. Cells were collected from 2 ml of culture by centrifugation at 4000 g for 10 min, washed once with PBS (phosphate-buffered saline; 137 mM NaCl, 2.7 mM KCl, 10 mM phosphate buffer, pH 7.4) and stored at −80 °C. The cell pellets were resuspended in 200 µl of B-PER bacterial protein extraction reagent (ThermoFisher Scientific) supplemented with 1 mg/ml lysozyme (Merck) and 0.1 mg/ml DNase I (Sigma) and lysed at room temperature for 15 min. Cell debris was removed by centrifugation (17,000 g, 5 min). An aliquot of cell lysate (50 µl) was mixed with 1 µl of 50 µM JFX549 and incubated for 30 min at 25 °C after which an equal volume of 2x Laemmli Sample Buffer (BIO-RAD) containing β-mercaptoethanol was added and samples were denatured at 98 °C for 5 min. For SDS-PAGE, 10 µl of each sample was loaded on a gel along with 1 µl of PageRuler Plus Prestained Protein Ladder were loaded on a Mini-Protean TGX gel (4-20%) and gel electrophoresis was performed according to the manufacturer’s protocol. JFX549-bound proteins were visualized using ChemiDoc MP imaging system. Band intensities were quantified using ImageJ. The gel was subsequently stained with InstantBlue Coomassie Protein Stain (ISB1L, Abcam) according to the manufacturer’s instructions and washed in water for 3 h. The total protein stained with Coomassie were visualized using ChemiDoc MP imaging system. Band intensities detected in the JFX549 fluorescence channel were normalized to prominent Coomassie-stained bands at ∼150–155 kDa, corresponding largely to the β and β′ subunits of RNA polymerase, used as an internal loading reference. Each experiment was performed three times. The mean HaloTag– Hsp15 signal in the WT HaloTag–Hsp15 strain was set to 1 and used as the reference for calculating relative abundance in all other samples. Error bars represent standard deviations between these independent experiments.

### BPA cross-linking

The *E. coli rpsG–HaloTag* strain carrying the plasmid pEVOL-pBpF (Addgene #31190) and either pQE-6His-Hsp15-124BPA or pQE-6His-Hsp15 was streaked onto LB agar plates supplemented with chloramphenicol (34 µg ml^−1^) and ampicillin (100 µg ml^−1^) and incubated overnight at 37 °C. Five to ten colonies were inoculated into 10 ml LB with chloramphenicol (34 µg ml^−1^) and ampicillin (100 µg ml^−1^) and grown to OD_600_ 0.6, followed by addition of 10 ml LB containing 20 mM arabinose, 1 mM IPTG, 2 mM Bpa (BLD Pharm), and the same antibiotics. Following induction, cells were grown for 4 h at 37 °C, after which erythromycin was added to a final concentration of 25 µg ml^−1^ and cultures were incubated for an additional 30 min. A 10 ml aliquot of each culture was irradiated with UV_365_ for 12 min at room temperature (UVP 2UV transilluminator, Analytik Jena), and the remaining 10 ml were maintained at room temperature without irradiation. Cells were cooled down in an ice bath and harvested by centrifugation (4000 g, 15 min, 4 °C).

The cell pellets were used for protein extraction using B-PER bacterial protein extraction reagent, HaloTag-fusions were labeled with JFX549 and visualized as described above for HaloTag-Hsp15. Cross-linking experiments were carried out in two independent replicates.

### Sample preparation for Cryo-EM

The *E. coli ΔssrA::kan* strain carrying a plasmid pQE-6His-Hsp15 plasmid was grown overnight in 10 ml LB supplemented with ampicillin (100 µg ml^−1^) and kanamycin (50 µg ml^−1^). An overnight culture (10 ml) was used to inoculate 2 L of 2×YT medium supplemented with ampicillin (100 µg ml^−1^). Cells were grown at 37 °C to an OD_600_ of 0.05, induced with IPTG (200 µM final concentration), and further cultured to an OD_600_ of 0.25. Erythromycin was then added to a final concentration of 25 µg ml^−1^ to induce ribosome collisions. Cultures were grown for an additional 30 min, reaching an OD_600_ of 0.5, and subsequently chilled on ice for 20 min. Cells were harvested by centrifugation at 4,000 × g for 15 min, washed once with ice-cold PBS, and stored at −80 °C.

Cell pellet was resuspended in 10 ml lysis buffer (20 mM HEPES (pH 7.6), 40 mM NH_4_Cl, 10 mM Mg(OAc)_2_, and 6 mM β-mercaptoethanol), supplemented with one tablet of cOmplete protease inhibitor and DNase I (0.1 mg ml^−1^). Cell suspension was disrupted by sonication (10 s on/30 s off, 6 cycles, repeated three times). Cell debris was removed by centrifugation at 18,000 × g for 30 min. The supernatant was filtered through a 0.45 µm filter and applied by gravity flow to a Co-NTA affinity column (1.5 ml resin) pre-equilibrated with lysis buffer. The column was washed twice with lysis buffer containing 10 mM imidazole (7 ml per wash), and bound proteins were eluted with lysis buffer supplemented with 250 mM imidazole (6 ml). Eluted fractions were concentrated using 10 kDa molecular weight cut-off centrifugal filters (Amicon) and buffer-exchanged into storage buffer (20 mM HEPES pH 7.6, 200 mM NH_4_Cl, 10 mM Mg(OAc)_2_, 6 mM β-mercaptoethanol added fresh) through repeated concentration and dilution cycles.

40 uL of ribosomes were sedimented through a 40-uL 50% sucrose cushion using an Optima MAX-XP ultracentrifuge with a TLA100 rotor (Beckman Coulter) at 80,000 RPM for 1.5 hrs. The pellet was re-suspended in 20 uL of polymix buffer (5 mM HEPES pH 7.5, 5 mM NH4Cl, 5 mM Mg(OAc)2, 100 mM KCl, 0.5 mM CaCl2, 8 mM putrescine, 1 mM spermidine, and 1 mM dithioerythritol) for a concentration of 140 nM and kept on ice until grid preparation.

### Cryo-EM

For single-particle cryo-EM analysis, a QuantiFoil 2/1 grid with 3 nm continuous carbon on 200 mesh Cu (Quantifoil Micro Tools GmbH) was glow-discharged at 0.4 mbar residual air with 20 mA current for 15 s. The grid was vitrified using a VitroBot mk IV (ThermoFisher Scientific) by applying 4 uL of sample to the carbon-side, incubated for 5 minutes, blotted for 5 s using Whatman 595 filter paper (Cytiva) and plunged into liquid ethane at -180 °C. The grid was screened and used for data collection on a Glacios 200 kV electron microscope equipped with a Selectris energy filter and a Falcon 4i direct electron detector (ThermoFisher Scientific). Data was collected using EPU version 3.14 (ThermoFisher Scientific) at 165,000x magnification (with a pixel-size in the sample plane of 0.686 Å), parallel illumination, energy-filter slit width 10 eV, dose-rate on camera over vacuum 8.58 e/pixel/s, total dose on sample 32.3 e/Å2, with a target defocus of -0.5 to -1.5 µm and 20 dose-fractions for a total of 57,327 exposures. Single-particle analysis was performed using CryoSPARC version v4.7.1^57^ (Supplementary Table S1). Movies were motion corrected using Patch Motion and optical parameters were estimated using Patch CTF. After curation, 46,454 exposures remained. Blob-picked particles (3.13 million) of diameter 220–280 Å were 3D classified using the heterogeneous refinement algorithm with 10 initial volumes generated by ab-inito reconstruction of both good and bad particles stemming from preliminary 2D classification. The particles were polished using reference-based motion correction and a final consensus reconstruction of 1.79 million particles at 2.11 Å was obtained by first performing homogeneous refinement with per-particle scale and defocus refinement, higher-order CTF parameter refinement (beamtilt, trefoil, CS and tetrafoil), magnification anisotropy refinement and Ewald Sphere curvature correction followed by separate beam-tilt and trefoil refinement for AFIS beam-shift groups. Global 3D classification without particle re-alignment into 10 classes filtered to 5 Å resulted in seven classical-state 70S classes (868,400 particles), one rotated 70S class (51,469) and three 50S classes (514,726, 172,595 and 163,930 particles). The 70S classical-state classes had varying levels of density for P-site tRNA and Hsp15. These were reconstructed into a 2.21-Å consensus 70S-classical-state reconstruction. After signal-subtraction of the 70S ribosome, focused 3D classification without particle re-alignment filtered to 6 Å within a mask for the Hsp15 and the P-site tRNA binding sites identified 10 out of 20 classes with significant Hsp15 density. The 443,984 particles in these classes were reconstructed into a 2.32 Å reconstruction and then subjected to 3D classification without particle re-alignment filtered to 6 Å within a mask for Hsp15, P-site tRNA and the SSU head. This identified four 70S classes with distinct Hsp15 density. Atomic modelling was done for the two 70S complexes (41,043 and 33,268 particles) and one 50S complex (514,726 particles) by first docking relevant parts of PDB 7K00, 8CGJ, 8CF1 into the maps. For Hsp15 and RsfS, an AlphaFold3 prediction^58^ was used as the starting point. P-site tRNAs for all three maps were modeled as tRNA^Asp^ (sequence obtained from GtRNAdb^59^). The models were interactively rebuilt and refined using Coot version 0.9.8.95^60^ and finally refined using Servalcat version 0.4.77^61^. Model validation was done using Molprobity^62^ and Phenix version 1.21.2^63^. Molecular figures were rendered using ChimeraX version 1.9^64^.

## Data availability

The fluorescence microscopy data generated in this study will be deposited in the publicly available SciLifeLab Repository. All unique biological materials will be available from the corresponding author upon request. Cryo-EM maps and models will be deposited in the Electron Microscopy Data Bank and the Protein Data Bank. Raw cryo-EM movies will be deposited in the EMPIAR database.

## Code availability

The computational code used for analysis and plotting will be available from the corresponding author and in the SciLifeLab Repository.

## ACKNOWLEDGEMENTS

The authors would like to thank the laboratory of Luke Lavis for providing the JFX549-HaloTag ligand. MATLAB scripts for figure generation were developed with assistance from ChatGPT and used exclusively for data visualization. This work was supported by the European Research Council (MJ: 947747-SMACK), the Swedish Research Council (MJ: 2023-03383, 2024-06104; MS: 2022-04511), Carl Tryggers Stiftelse för Vetenskaplig Forskning (MJ, ACS: 17:226), and The Wenner-Gren Foundations (MJ, ACS). The computations and data management were enabled by resources provided by the Swedish National Infrastructure for Computing at UPPMAX, partially funded by the Swedish Research Council through grant agreement no. 2018-05973. Cryo-EM grid preparation and data collection was done at the Cryo-EM Uppsala facility, funded by the Department of Cell and Molecular Biology, the Disciplinary Domains of Science and Technology and of Medicine and Pharmacy at Uppsala University.

