## Supplementary Material for "Ribosome collisions trigger tmRNA-mediated rescue through mRNA disengagement"

[*] *These authors contributed equally*

**Supplementary note 1. Rationale for coarse-graining in HMM analysis.**

Hidden Markov modelling of single-molecule tracking data for ribosomal subunits and associated factors reveals that their diffusion is more accurately described by broad coefficient distributions than by sharply separated states[1–4]. This complexity likely reflects the diverse biological contexts in which ribosomes exist, ranging from rapidly diffusing mRNA-free subunits to polysomes with varying ribosome occupancy per mRNA, and to ribosomes further constrained through association with the nucleoid via RNA polymerase or with the membrane via translocons, resulting in progressively slower diffusion. Importantly, this complexity is not specific to HMM analysis but is also evident in simpler fits of diffusion step-length histograms, which are better described by multiple states than by a single-state model[4]. However, while models with a larger number of states provide a better statistical description of the data, they are not necessarily optimal for biological interpretation. We therefore model the data with larger HMM models to capture the full diffusion landscape and subsequently coarse-grain the resulting states into biologically interpretable classes. This strategy reduces the risk of overlooking low-populated states. In addition, introducing multiple diffusive states prior to coarse-graining compensates for the exponential dwell-time assumption inherent to HMM, allowing non-exponential residence times to be effectively represented.

In this work, HMM models containing 3 to 8 states produced comparable results after coarse-graining for all factors analyzed. For simplicity of presentation, we therefore use coarse-grained 3-state models throughout the manuscript.

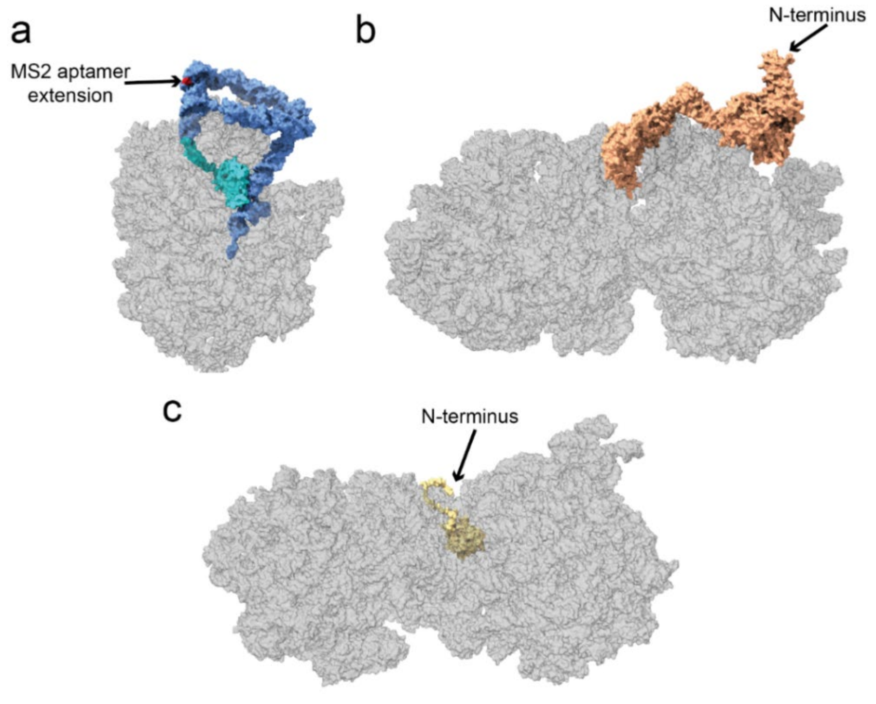

**Supplementary Fig. 1. Cryo-EM structures of tmRNA-, HrpA- and SmrB-bound ribosomes highlighting labelling sites. a.** Cryo-EM structure of an accommodated trans-translation complex on a stalled *E. coli* ribosome (PDB 7AC7[5]). tmRNA is shown in blue and SmpB in cyan. The position nucleotide 175, used for insertion of the MS2 aptamer, is highlighted in red and has been shown to tolerate incorporation of heterologous sequences[6]. **b**. Cryo-EM structure of an *E. coli* disome bound by HrpA (PDB 9GFT). HrpA is shown in orange, and the position of the N terminus used for HaloTag fusion is indicated. **c**. Cryo-EM structure of an *E. coli* disome bound by SmrB (PDB 7QGN, stalled 70S; PDB 7QGR, collided 70S). SmrB is shown in yellow, and the position of the N terminus used for HaloTag fusion is indicated.

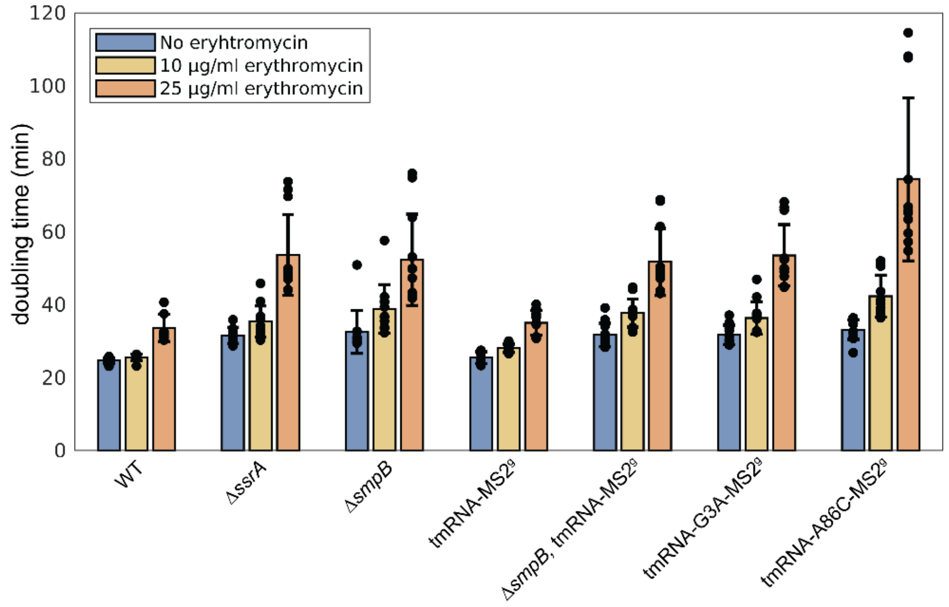

**Supplementary Fig. 2. Doubling times of WT strain and trans-translation mutants in the presence and absence of erythromycin.** Bars represent averages from independent experiments, and error bars indicate standard deviations between experiments.

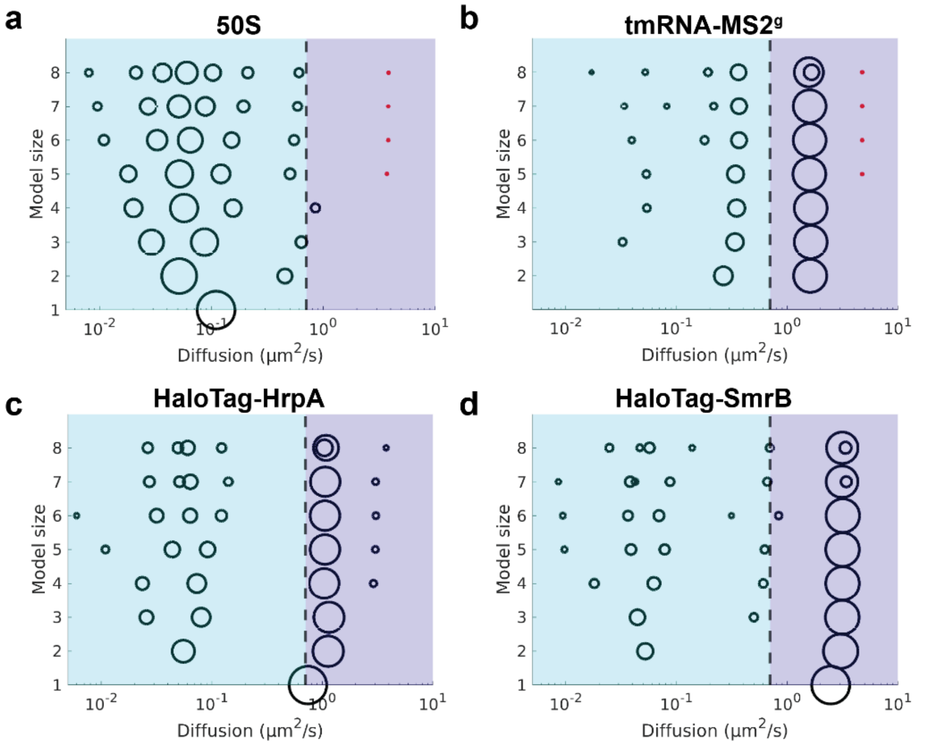

**Supplementary Fig. 3. Fitted HMM models of diffusion states for 50S, tmRNA-MS2$^{g}$, HaloTag-HrpA, and HaloTag-SmrB.** The area of each circle represents the relative occupancy of the corresponding diffusion state for each model size. Diffusion states with occupancies below 1% are marked with red asterisks. Black dashed line indicates the threshold at 0.7 µm$^2$ s$^{-1}$ used to distinguish ribosome-bound states (cyan area) from free factors (violet area). The data show results from HMM fitting of datasets from all individual experiments combined.

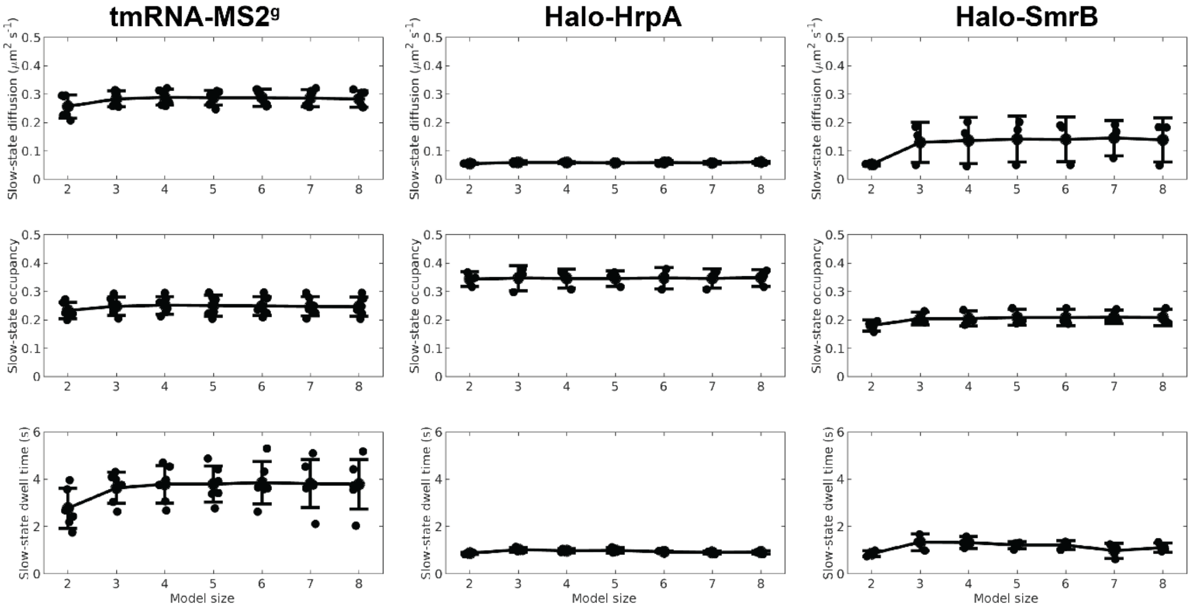

**Supplementary Fig. 4. Slow diffusion state parameters across different HMM model sizes after coarse-graining.** Occupancies, dwell times and diffusion coefficients of tmRNA–MS2$^{g}$, HaloTag–HrpA and HaloTag–SmrB in the slow diffusion state across different HMM model sizes. Data are shown after coarse-graining using a 0.7 μm² s$^{-1}$ threshold to define ribosome-bound states. Dots represent values from independent experiments, lines represent averages of all individual experiments, and error bars indicate standard deviations between experiments.

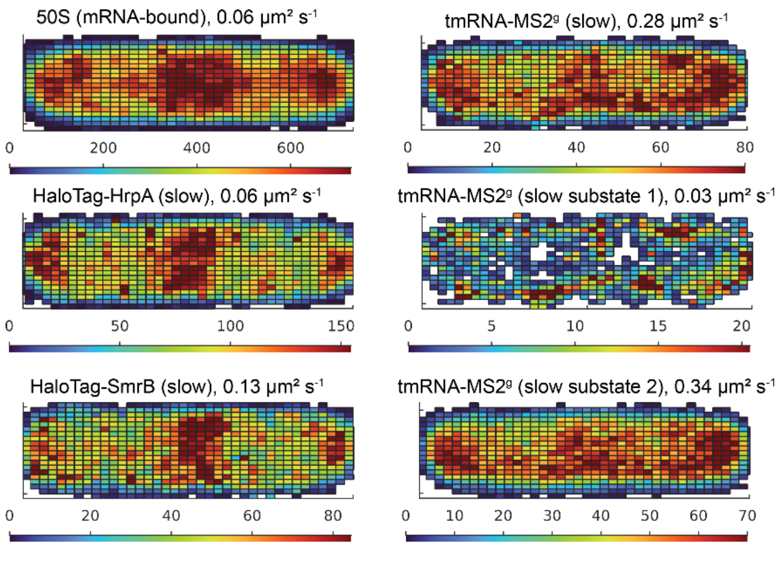

**Supplementary Fig. 5. Spatial distribution of the slow diffusion states for tmRNA-MS2$^{g}$, HaloTag-HrpA, HaloTag-SmrB, and mRNA-bound 50S.** Distributions for mRNA-bound 50S were obtained from a three-state HMM model coarse-grained by grouping states with diffusion coefficients below 0.25 μm² s$^{-1}$, with the threshold defined according to our previous work[1]. Distributions for the slow states of HrpA–HaloTag, SmrB–HaloTag and tmRNA–MS2$^{g}$ were obtained from three-state HMM models coarse-grained by grouping states with diffusion coefficients below 0.7 μm² s$^{-1}$. Distributions for tmRNA–MS2$^{g}$ slow substates 1 and 2 were derived from three-state HMM models, with the first and second states shown separately. The average diffusion coefficients in corresponding states are indicated.

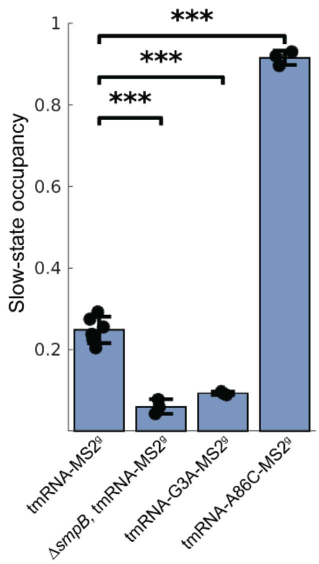

**Supplementary Fig. 6. Effect of tmRNA mutations and *smpB* deletion on ribosome-bound state occupancy of tmRNA-MS2$^{g}$.** Ribosome-bound state occupancies for tmRNA-MS2$^{g}$, tmRNA-G3A-MS2$^{g}$, tmRNA-A86C-MS2$^{g}$ in WT strain and tmRNA-MS2$^{g}$ in *ΔsmbB* strain. The data are derived from coarse-grained 3-state HMM models. Bars represent averages from independent experiments, and error bars indicate standard deviations between experiments. Statistical significance between groups was assessed using a two-sided unpaired t-test. P-values are indicated in the figure as follows: P < 0.001 (***).

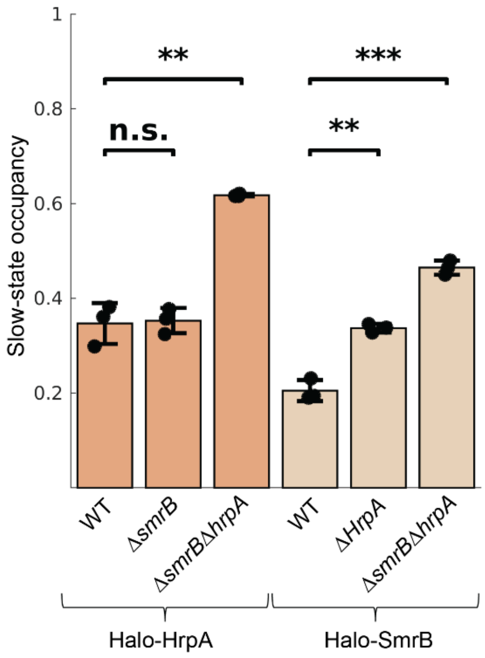

**Supplementary Fig. 7. Effect of *smrB* and *hrpA* deletion on ribosome-bound state occupancy of HaloTag–HrpA and HaloTag–SmrB.** The data are derived from coarse-grained 3-state HMM models. Bars represent averages from independent experiments, and error bars indicate standard deviations between experiments. Statistical significance between groups was assessed using a two-sided unpaired t-test. P-values are indicated in the figure as follows: P < 0.01 (**), P < 0.001 (***), and not significant (n.s.) for P ≥ 0.05.

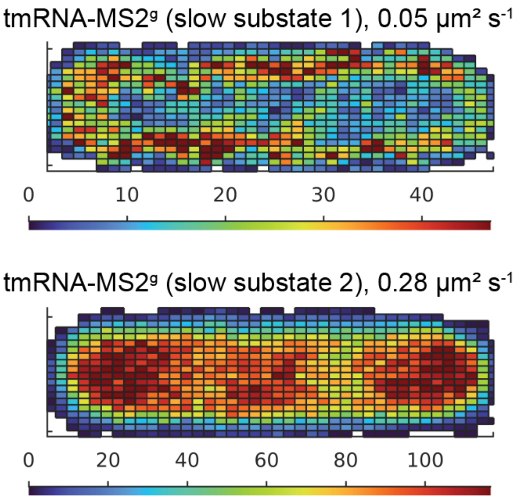

**Supplementary Fig. 8. Spatial distribution of the slow diffusion substates of tmRNA-MS2g in erythromycin treated cells.** Distributions were derived from a three-state HMM model. The average diffusion coefficients in those states are indicated.

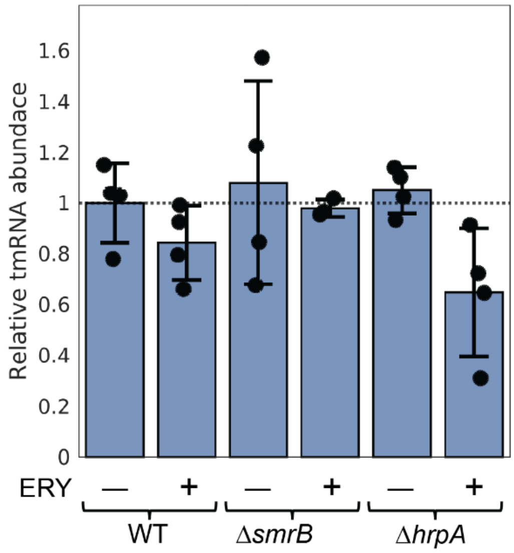

**Supplementary Fig. 9. Relative tmRNA abundance in WT, *ΔsmrB* and *ΔhrpA* strains in the absence and presence of erythromycin.** Bars represent averages from independent experiments, and error bars indicate standard deviations between experiments. Although reduced tmRNA levels in the *ΔhrpA* strain would be expected to increase ribosome-bound state occupancy under unchanged usage, we instead observe a decrease (Fig. 2c).

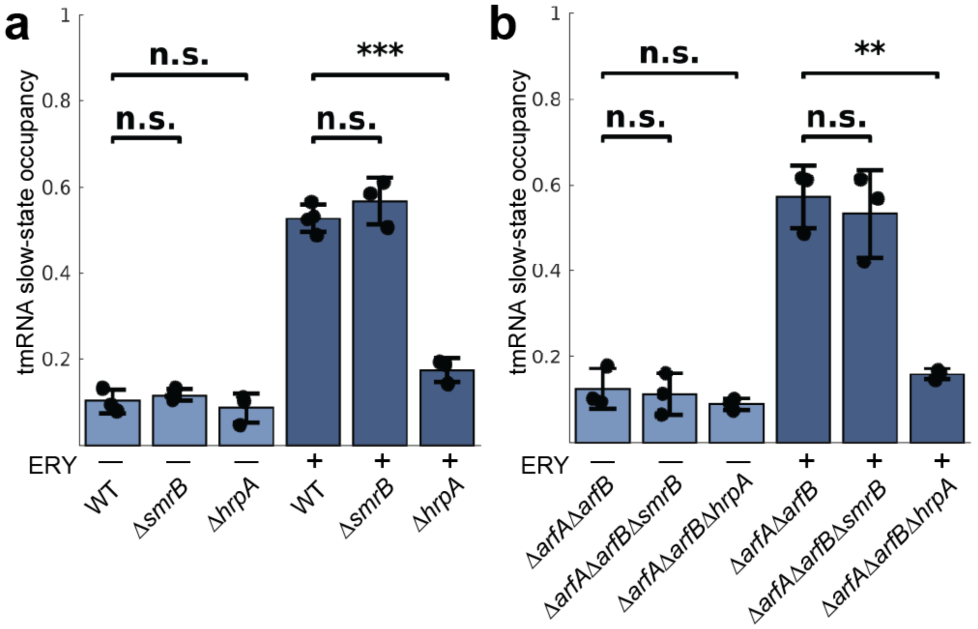

**Supplementary Fig. 10. Effect of *smrB* and *hrpA* deletions on recruitment of tmRNA-MS2$^{p}$ to ribosomes. a.** Ribosome-bound state occupancies of tmRNA-MS2$^{p}$ in WT, *ΔsmrB* and *ΔhrpA* strains without and with erythromycin treatment. **b.** Ribosome-bound state occupancies of tmRNA-MS2$^{p}$ in *ΔarfAΔarfB*, *ΔarfAΔarfBΔsmrB* and *ΔarfAΔarfBΔhrpA* strains without and with erythromycin treatment. Data are derived from coarse-grained 3-state HMM models. Bars represent averages from independent experiments, and error bars indicate standard deviations between experiments. Statistical significance between groups was assessed using a two-sided unpaired t-test. P-values are indicated in the figure as follows: $P < 0.01$ (**), $P < 0.001$ (***), and not significant (n.s.) for $P \geq 0.05$.

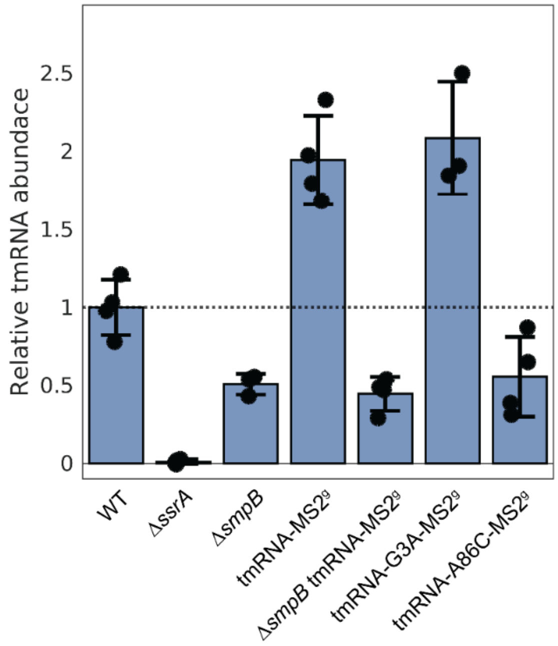

**Supplementary Fig. 11. Relative tmRNA abundance in WT and tmRNA-MS2$^{g}$ strains.** Bars represent averages from independent experiments, and error bars indicate standard deviations between experiments.

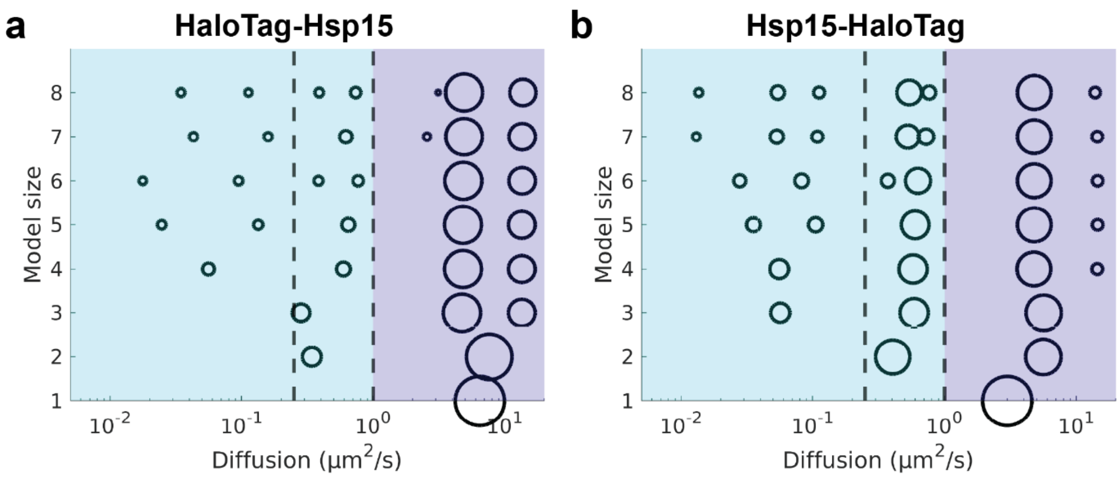

**Supplementary Fig. 12. Fitted HMM diffusion models for HaloTag–Hsp15 and Hsp15–HaloTag.** The area of each circle represents the relative occupancy of the corresponding diffusion state. A black dashed line at 1 μm² s⁻¹ indicates the threshold used to distinguish ribosome-bound states (cyan region) from freely diffusing factors (violet region). A black dashed line at 0.25 μm² s⁻¹ separates membrane-associated and cytosolic Hsp15 populations. The data show results from HMM models for combined datasets.

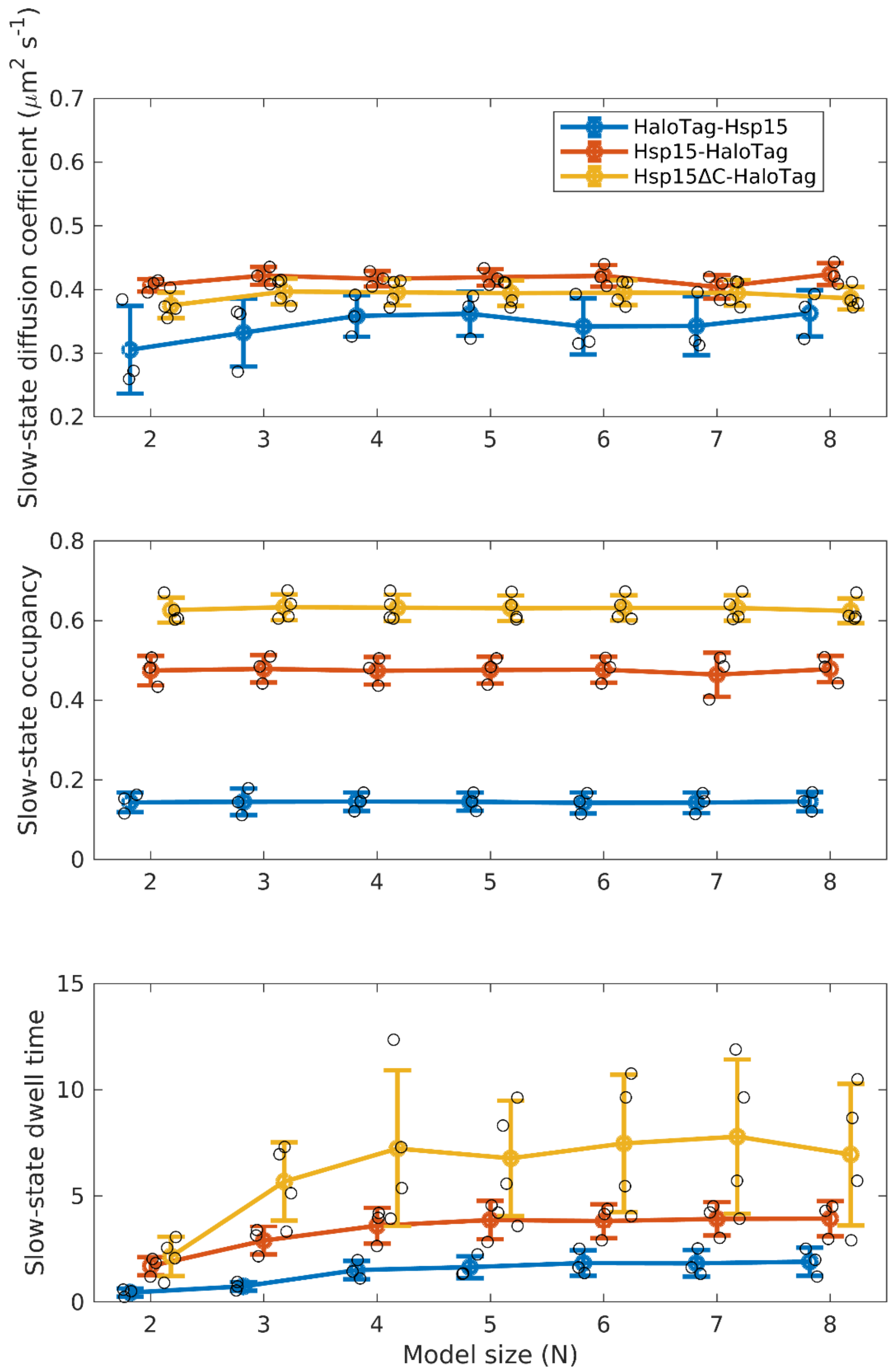

**Supplementary Fig. 13. Hsp15 slow diffusion state parameters across HMM model sizes after coarse-graining.** Occupancies, dwell times and diffusion coefficients of HaloTag-Hsp15, Hsp15-HaloTag and Hsp15ΔC-HaloTag mutant in the slow diffusion state across different HMM model sizes. Data are shown after coarse-graining using a 1 μm² s⁻¹ threshold to define ribosome-bound states. Values represent averages from independent experiments, and error bars indicate standard deviations between experiments.

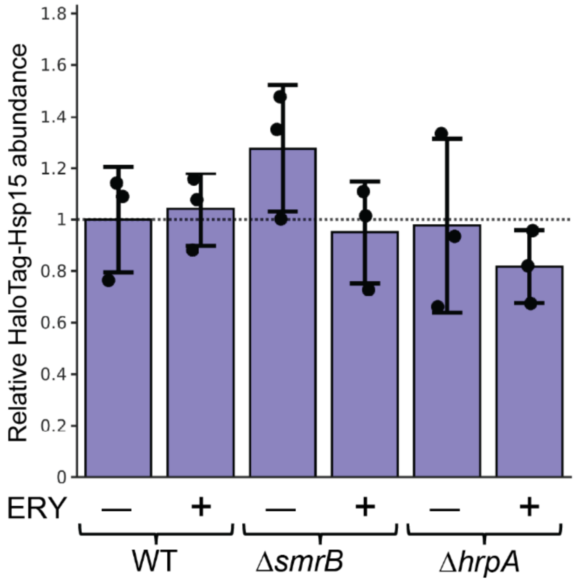

**Supplementary Fig. 14. Relative abundance of HaloTag-Hsp15 in WT, *ΔsmrB* and *ΔhrpA* strains in the absence and presence of erythromycin.** Bars represent averages from independent experiments, and error bars indicate standard deviations between experiments.

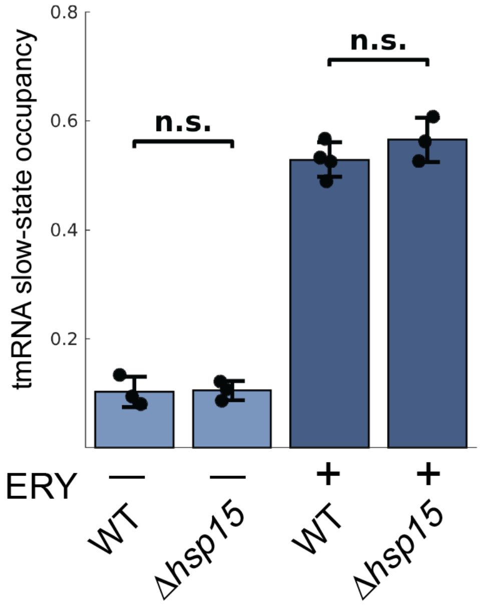

**Supplementary Fig. 15. Ribosome-bound occupancies of tmRNA-MS2$^p$ in WT and *Δhsp15* strains.** Data are derived from coarse-grained 3-state HMM models. Bars represent averages from independent experiments, and error bars indicate standard deviations between experiments. Statistical significance between groups was assessed using a two-sided unpaired t-test. P-values are indicated in the figure as follows: not significant (n.s.) for $P \geq 0.05$.

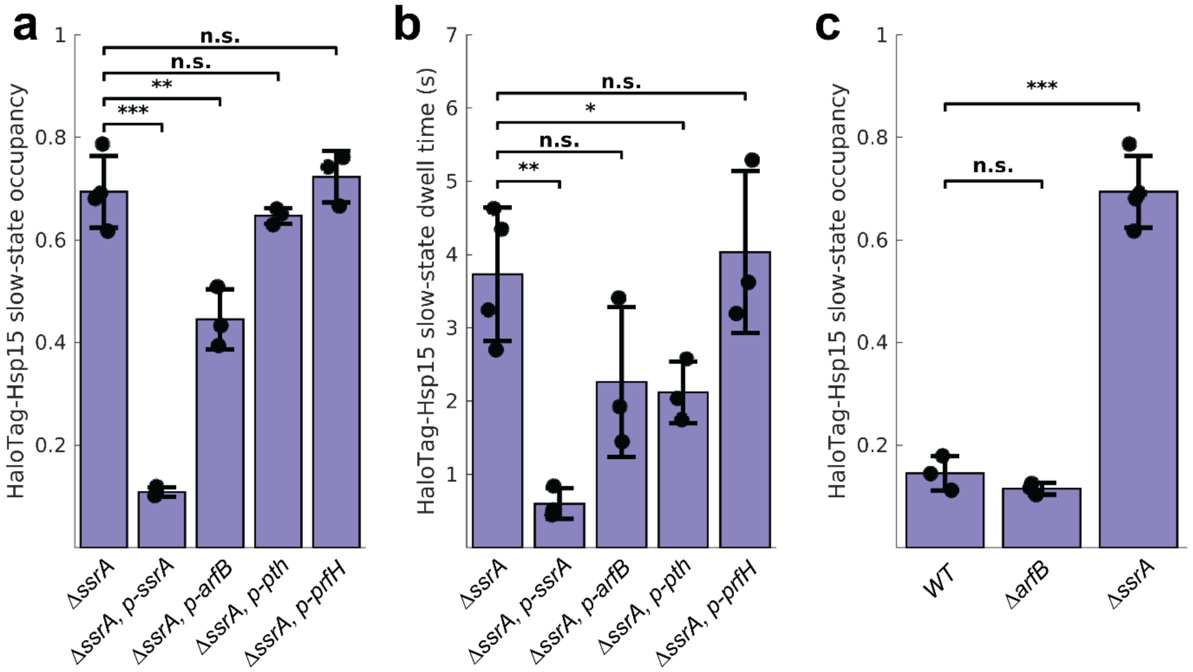

**Supplementary Fig. 16. Effect of Pth, PrfH and ArfB overexpression and *arfB* deletion on association of HaloTag–Hsp15 with ribosomes. a-b.** Ribosome-bound state occupancies (**a**) and dwell times (**b**) of HaloTag–Hsp15 in the *ΔssrA* strain carrying plasmids expressing *ssrA*, *arfB*, *pth* or *prfH*. **c**. Ribosome-bound state occupancies of HaloTag–Hsp15 in the WT, *ΔarfB, and ΔssrA* strains. Data are derived from coarse-grained 3-state HMM models. Bars represent averages from independent experiments, and error bars indicate standard deviations between experiments. Statistical significance between groups was assessed using a two-sided unpaired t-test. P-values are indicated in the figure as follows: $P < 0.05$ (*), $P < 0.01$ (**), $P < 0.001$ (***), and not significant (n.s.) otherwise.

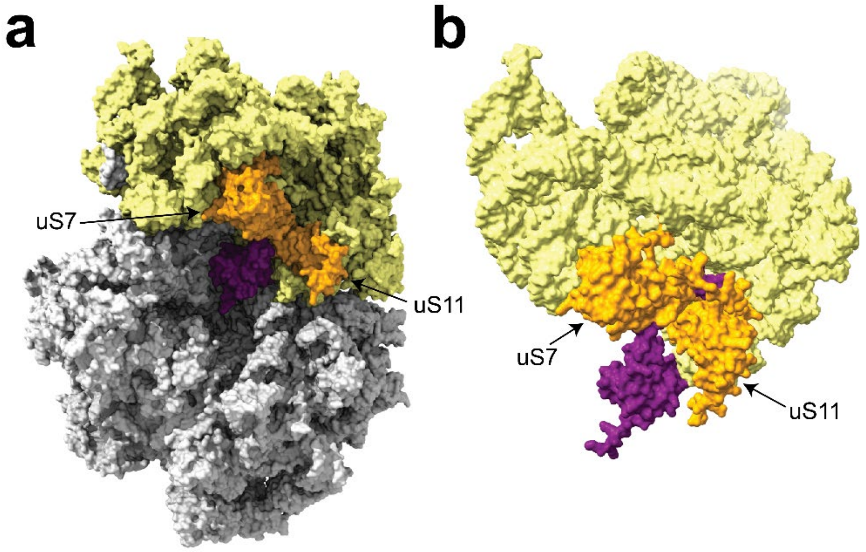

**Supplementary Fig. 17. Prediction of Hsp15 positioning on the 70S ribosome. a**. The previously resolved structure of the 70S ribosome (PDB 7K00) was overlaid with the 50S–peptidyl-tRNA–Hsp15 complex (PDB 8AP4) to predict potential interactions with the 30S subunit. The unresolved C-terminal α-helix of Hsp15 is predicted to extend into the E-site mRNA channel near uS7 and uS11 and would likely sterically clash with mRNA in this channel. **b**. AlphaFold modelling of *E. coli* Hsp15 in the context of 16S rRNA, uS7, and uS11 predicts that the C-terminal α-helix occupies the E-site mRNA channel and is positioned at the interface between 16S rRNA, uS7, and uS11. 50S is shown in grey, 30S in **a** and 16S rRNA in **b** are shown in yellow, uS7 and uS11 are shown in orange, Hsp15 is shown in violet.

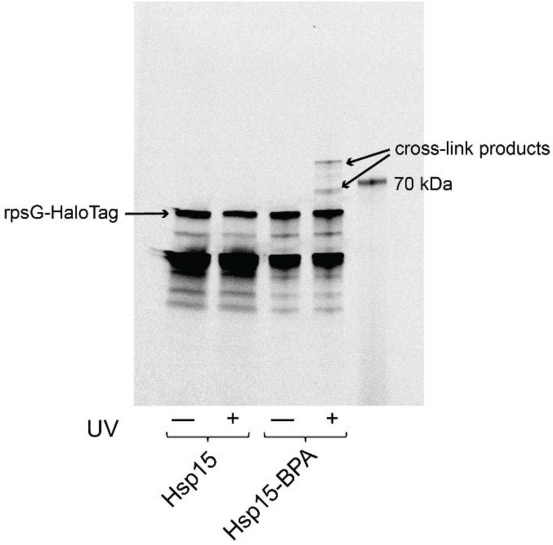

**Supplementary Fig. 18. SDS–PAGE analysis of uS7–HaloTag cross-linked to Hsp15 containing the photoinducible amino acid p-benzoyl-L-phenylalanine.** 6His–Hsp15 or 6His–Hsp15–Bpa was overexpressed in Escherichia coli carrying a chromosomal C-terminal HaloTag fusion to uS7 (rpsG) and the pEVOL-pBpF plasmid. Following induction, cell cultures were either exposed to UV irradiation to induce cross-linking or left untreated. Cell lysates were labelled with JFX549 to detect the uS7–HaloTag fusion and analyzed by SDS–PAGE. The experiment was independently repeated twice.

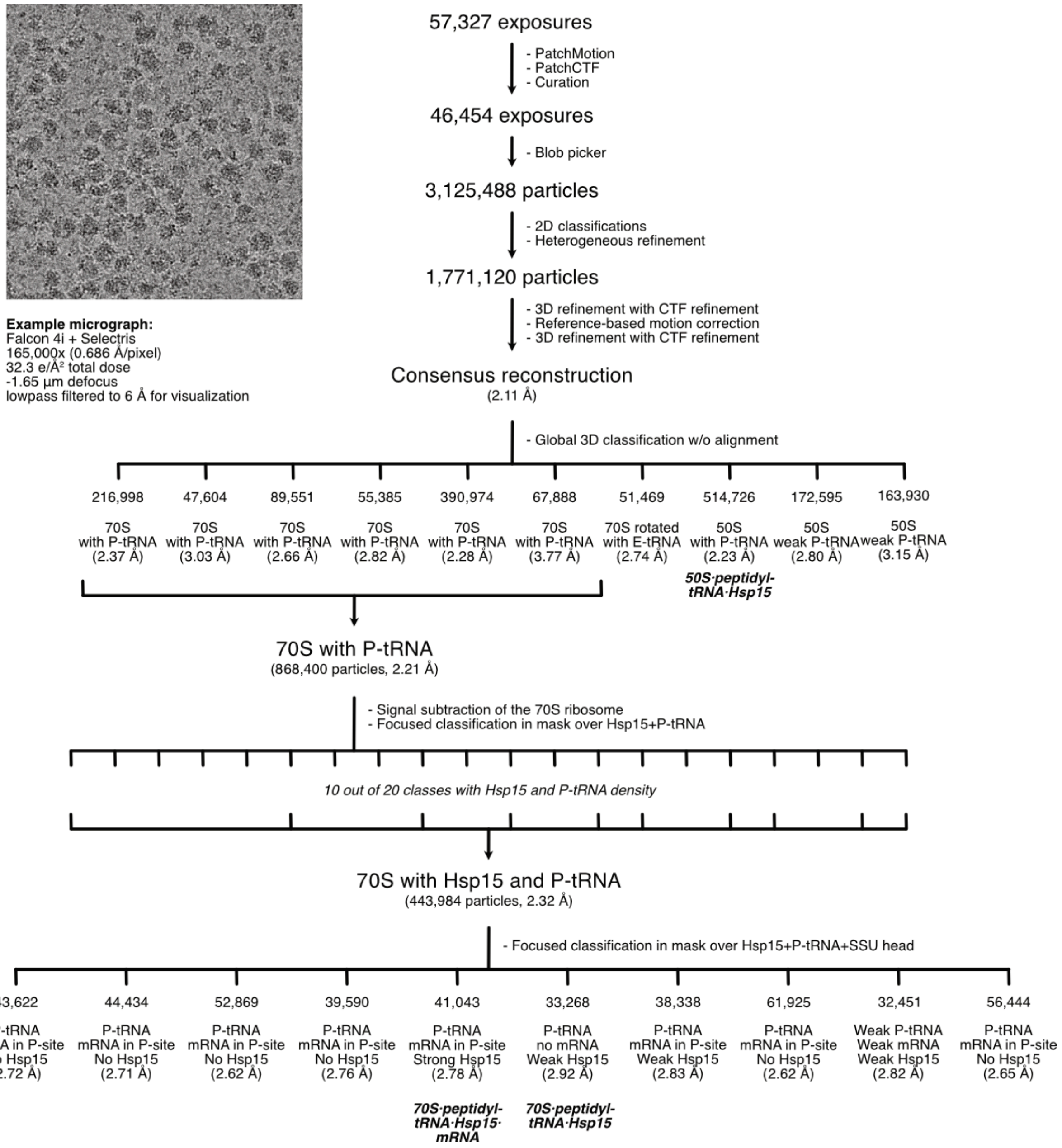

**Supplementary Fig. 19. Cryo-EM single-particle analysis workflow.** Representative motion corrected and low pass filtered (6 Å) micrograph shown.

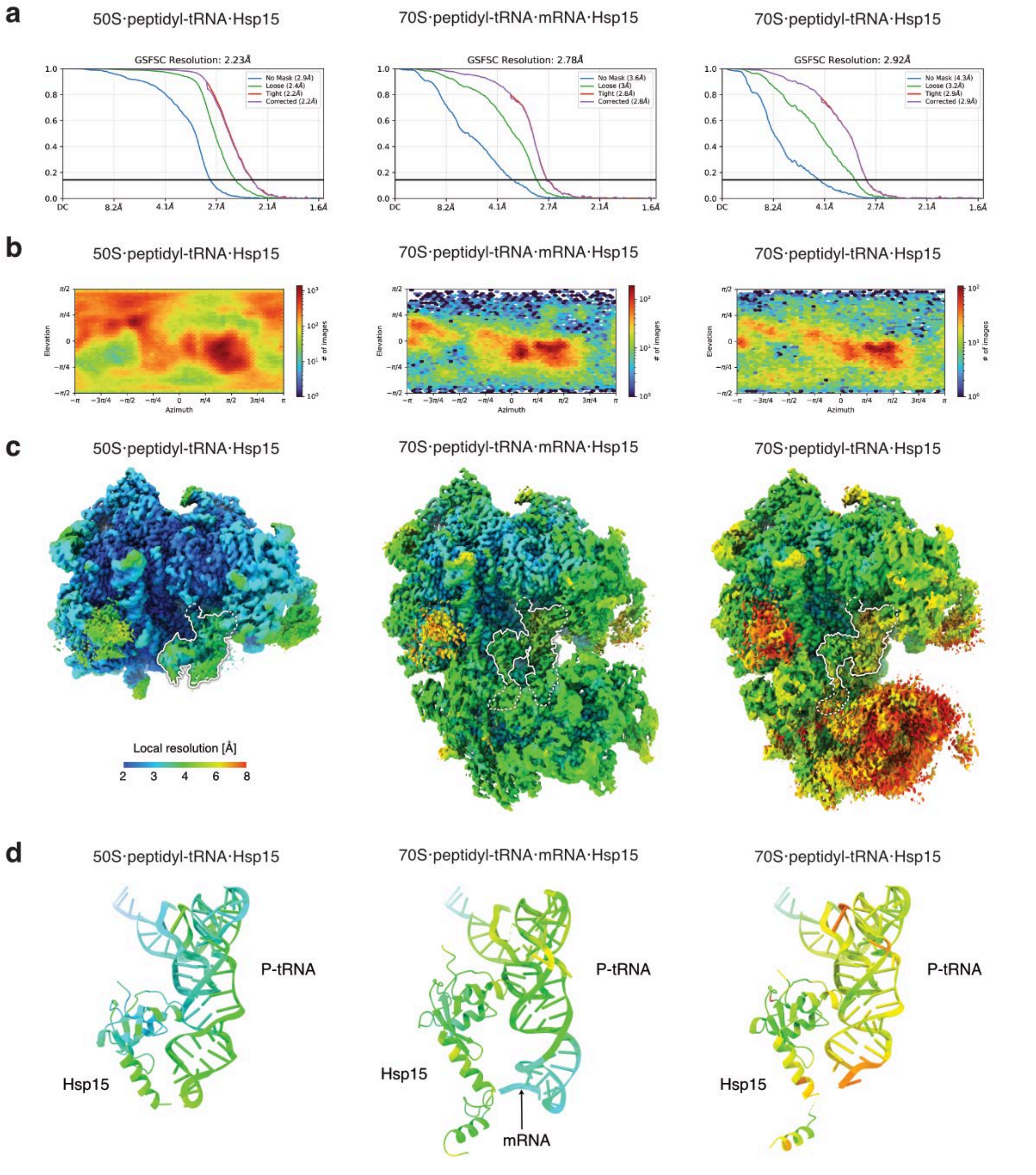

**Supplementary Fig. 20. Cryo-EM reconstruction resolution estimations. a.** Gold-standard Fourier Shell Correlation plots from final CryoSPARC homogeneous refinement jobs. **b.** Euler angle distribution plots from final CryoSPARC homogeneous refinement jobs. **c.** Local resolution estimation using CryoSPARC local resolution job with default parameters used to color the reconstruction map using ChimeraX with the palette blue for 2 Å, cyan for 3 Å, light green for 4 Å, yellow for 6 Å and red for 8 Å. **d.** Refined models for Hsp15, P-tRNA and mRNA shown as cartoons and colored according to the estimated local resolution as the average value per reside using the same palette as in **c**.

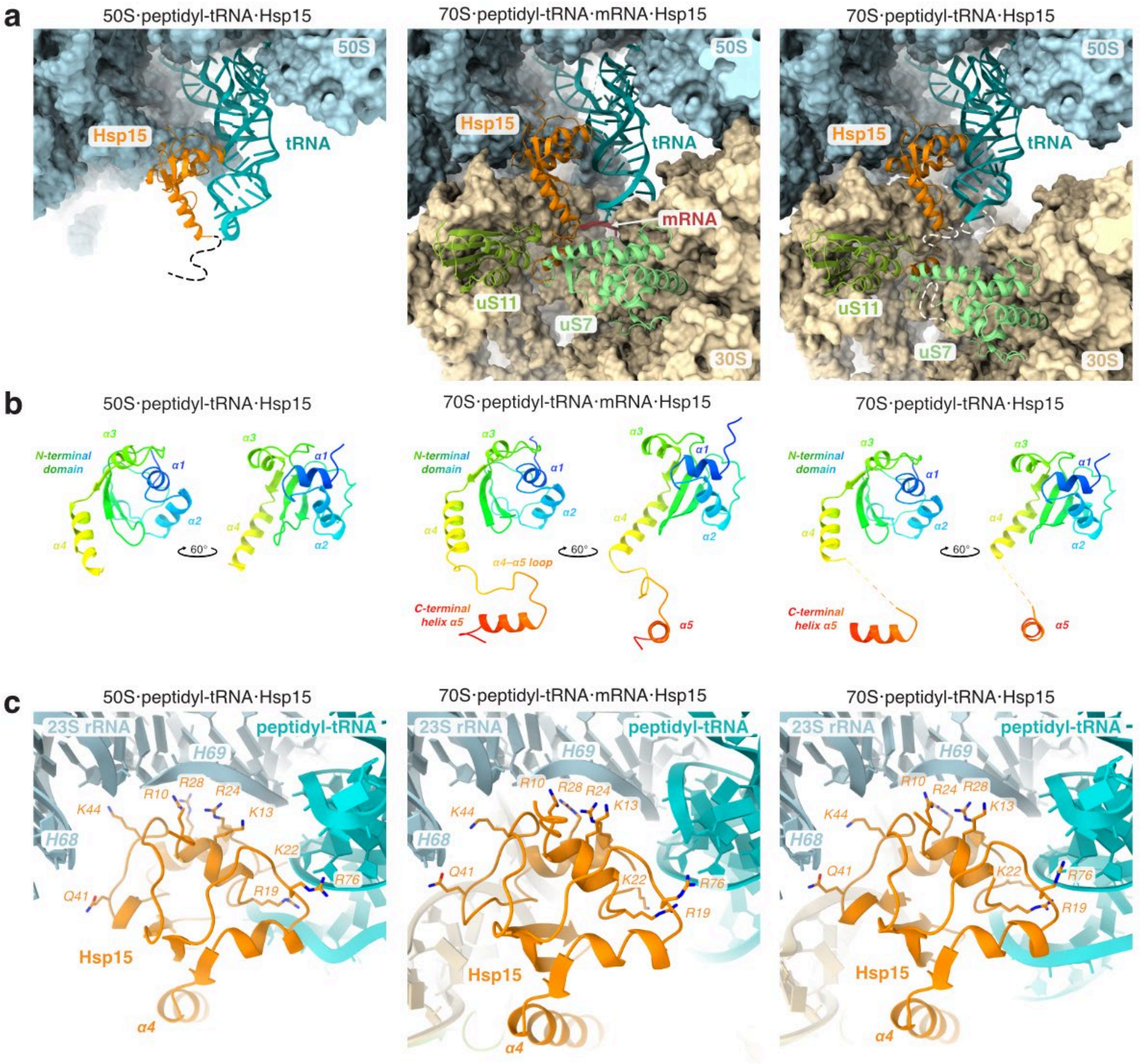

**Supplementary Fig. 21. Hsp15 conformation in ribosomal complexes. a.** Position of Hsp15 (orange) relative to peptidyl-tRNA (teal), uS7 (mint green), uS11 (yellow green), mRNA (red), 50S (light blue surface), 30S (wheat colored surface) for 50S complex (left), 70S complex with mRNA (middle) and 70S complex without mRNA (right). Flexible loops indicated with dashed lines. **b.** Conformation of Hsp15 in the three complexes. Rainbow coloring from N-terminus in blue to C-terminus in red. **c.** Polar contacts by the N-terminal domain to the 23S rRNA and the peptidyl-tRNA.

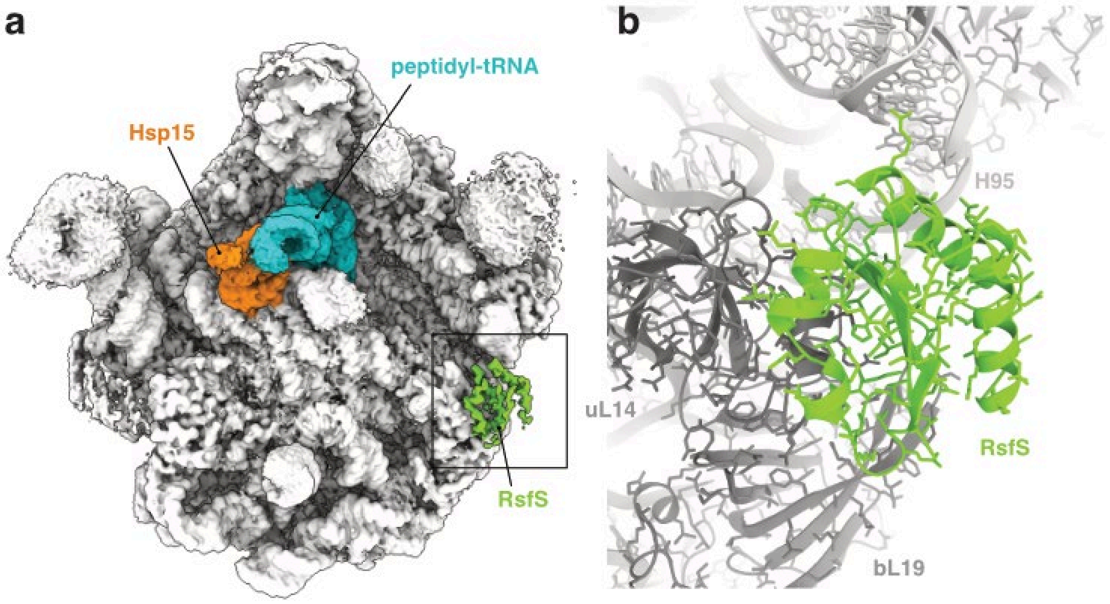

**Supplementary Fig. 22. Hibernation factor RsfS bound to the 50S·peptidyl-tRNA·Hsp15 complex. a.** Crown view with density for partial occupancy of RsfS bound to the LSU. Hsp15 orange, peptidyl-tRNA teal, 50S ribosome white, RsfS green. **b.** Close-up showing the refined model of RsfS (green) bound to r-proteins uL14 and bL19 (both dark gray) and to 23S rRNA (light gray) helix H95.

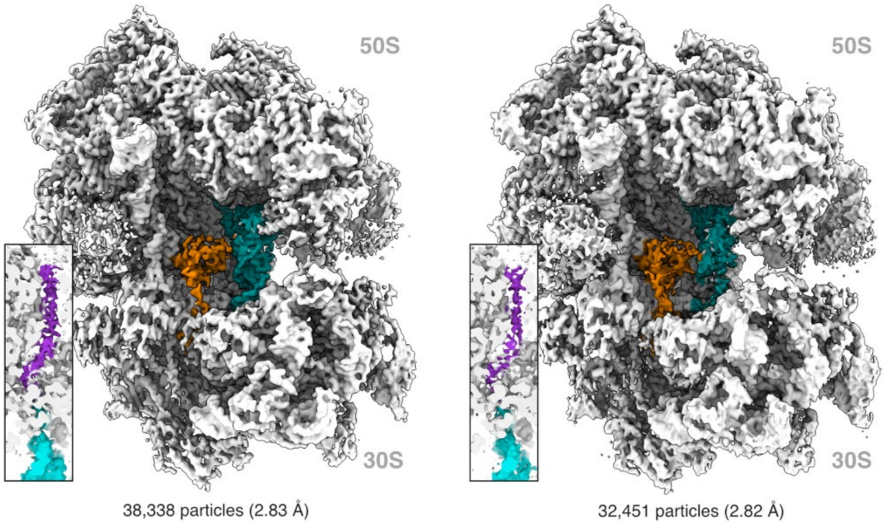

**Supplementary Fig. 23. Additional 70S complexes with Hsp15, peptidyl-tRNA and mRNA.** Same view as Fig 4a.

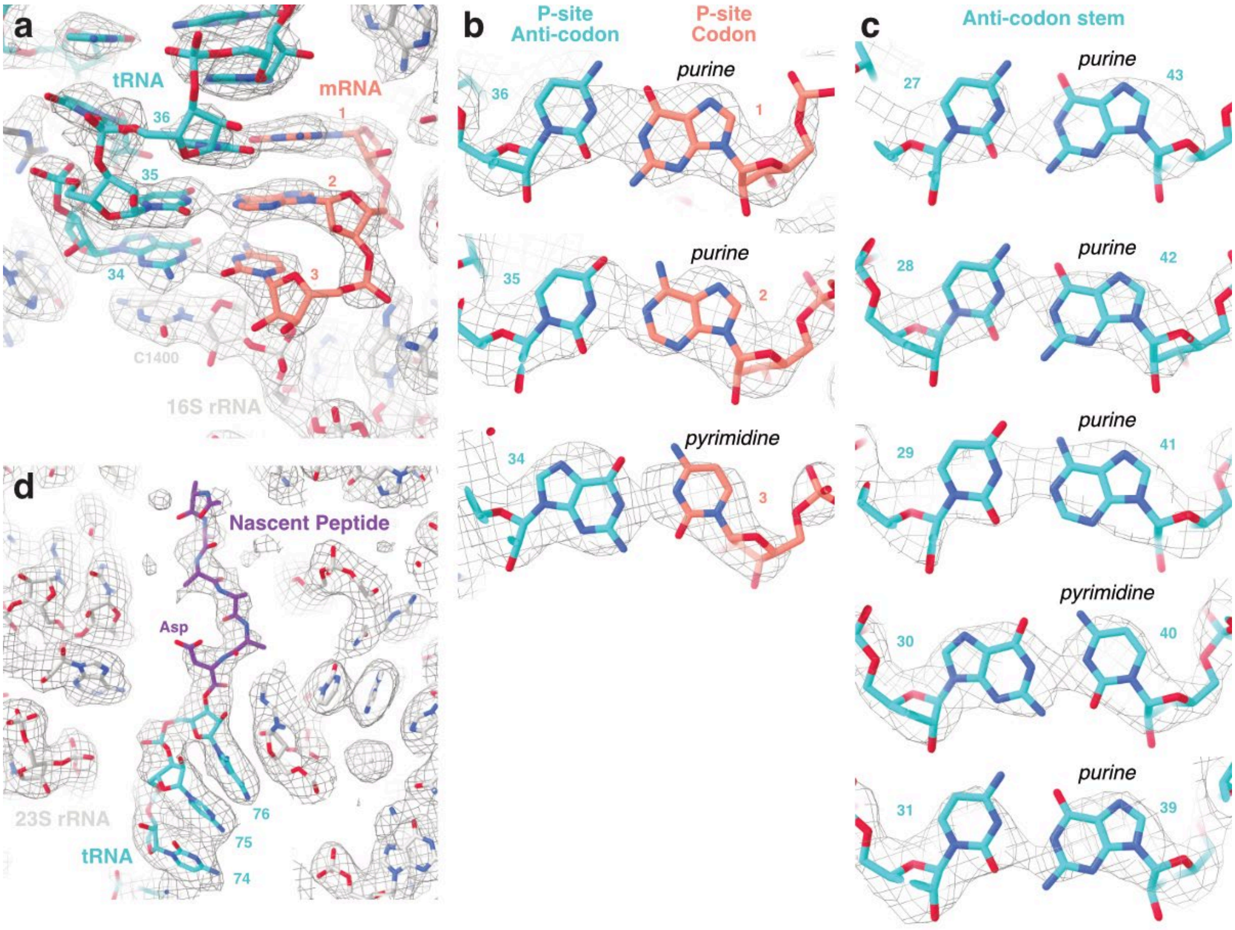

**Supplementary Fig. 24. Identification of the peptidyl-tRNA in complex 70S·peptidyl-tRNA·mRNA·Hsp15. a.** mRNA-tRNA codon-anticodon mini-helix at the SSU P-site. Numbers are residue numbers for the tRNA (cyan) and positions for the P-site mRNA (pink). Cryo-EM map shown as mesh. **b.** Close-up of each codon-anticodon base-pair with density shown as mesh to support the assignment of purines and pyrimidines. **c.** Close-up of each base-pair in the tRNA anticodon stem with density shown as mesh to support the assignment of purines and pyrimidines with residue numbers indicated. **d.** CCA-end of the tRNA molecule with one possible way to model the nascent polypeptide chain in purple (the peptide is not included in the deposited model).

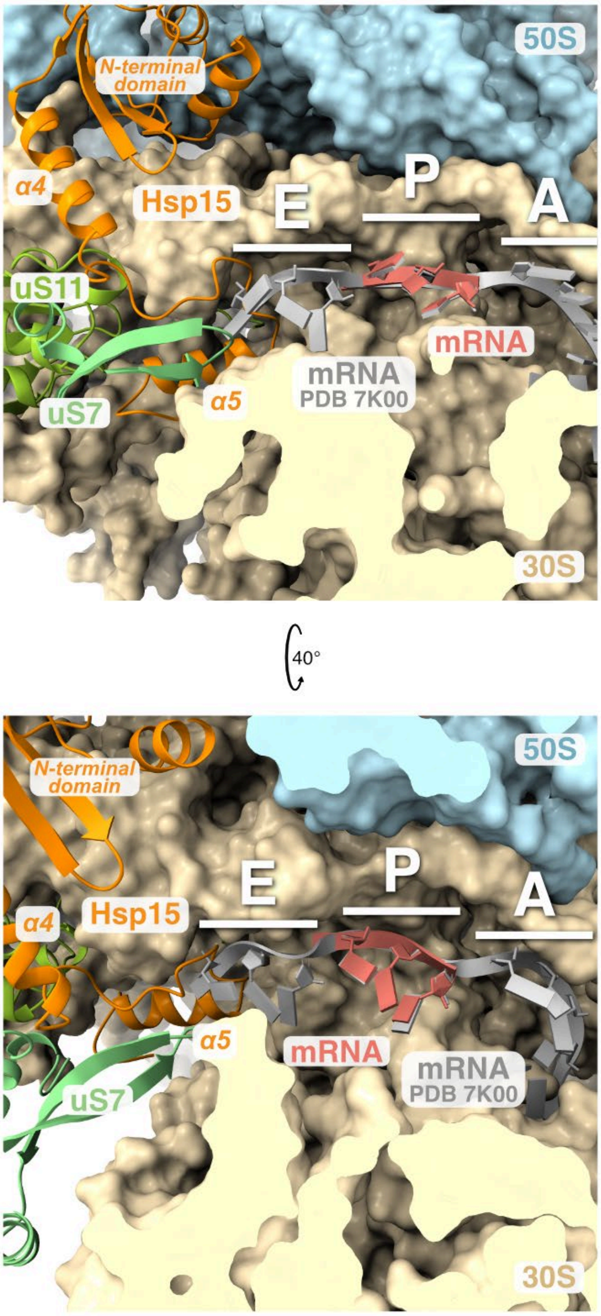

**Supplementary Fig. 25. Obstruction of the mRNA channel by the C-terminus of Hsp15.** The 70S·peptidyl-tRNA·mRNA·Hsp15 complex shown overlayed with the mRNA from PDB 7K00 (shown in grey). The α4–α5 loop and the C-terminal a5-helix of Hsp15 (orange) blocks the mRNA channel at the E site. 50S light blue surface, 30S yellow surface, uS7 mint green, mRNA red.

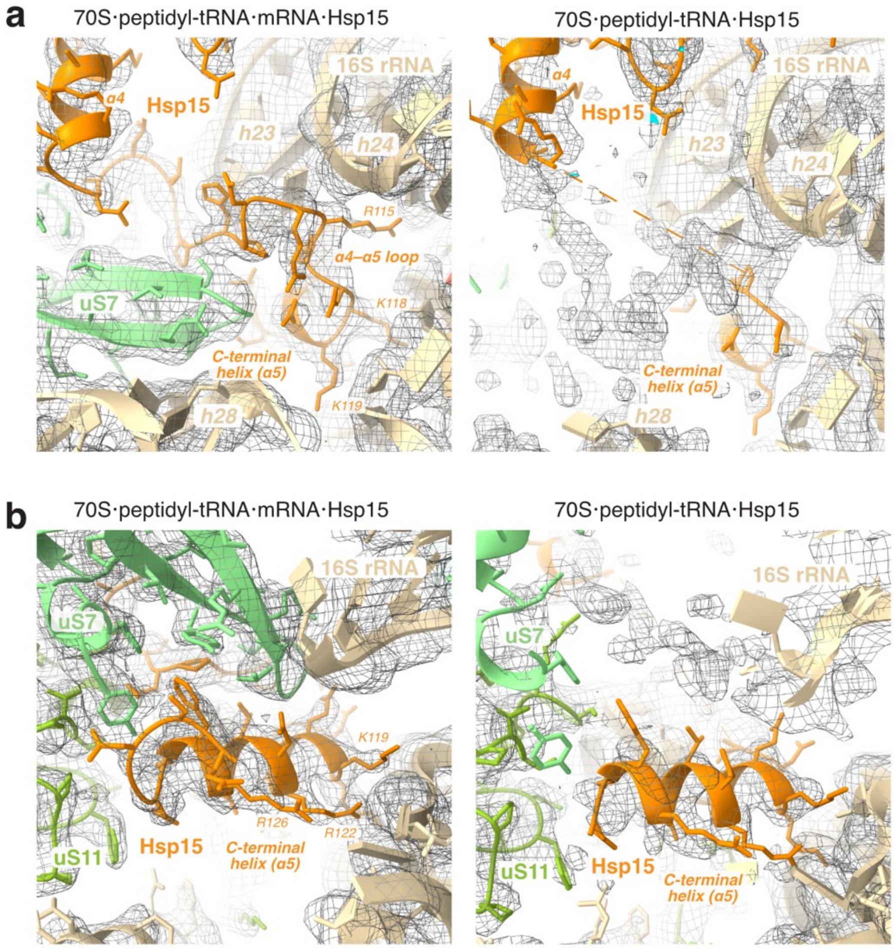

**Supplementary Fig. 26. Comparison of the C-terminal region of Hsp15 in the two 70S complexes. a.** α4–α5 loop. Same view as Fig 4e. **b.** C-terminal helix. Same view as Fig 4f.

**Supplementary Table S1. Cryo-EM data collection, refinement and validation statistics.**

| | #1 50S·peptidyl-tRNA·Hsp15 (EMDB-xxxx) (PDB xxxx) | #2 70S·peptidyl-tRNA·mRNA·Hsp15 (EMDB-xxxx) (PDB xxxx) | #3 70S·peptidyl-tRNA·Hsp15 (EMDB-xxxx) (PDB xxxx) |
|---|---|---|---|
| **Data collection and processing** | | | |
| Magnification | 165,000x | 165,000x | 165,000x |
| Voltage (kV) | 200 | 200 | 200 |
| Electron exposure (e–/Å$^2$) | 32.3 | 32.3 | 32.3 |
| Defocus range (μm) | -0.5 to -1.5 | -0.5 to -1.5 | -0.5 to -1.5 |
| Pixel size (Å) | 0.686 | 0.686 | 0.686 |
| Symmetry imposed | C1 | C1 | C1 |
| Initial particle images (no.) | 3,125,488 | 3,125,488 | 3,125,488 |
| Final particle images (no.) | 514,726 | 41,043 | 33,268 |
| Map resolution (Å) | 2.23 | 2.78 | 2.92 |
| FSC threshold | 0.143 | 0.143 | 0.143 |
| Map resolution range (Å) | 1.75–38 | 1.72–47 | 1.72–47 |
